# KIF1A-driven entry of degradative lysosomes into the axon is developmentally regulated by the microtubule environment at the axon initial segment

**DOI:** 10.64898/2026.10.29.762862

**Authors:** Nireekshit Addanki Tirumala, Surbhi Verma, Juan S. Bonifacino

## Abstract

Neuronal function requires precise transport of lysosomal vesicles along highly polarized processes, yet how this trafficking is spatially and developmentally regulated remains poorly understood. Here, we show that the neuronally enriched kinesin-3 motor KIF1A is required for dendritic and axonal localization of lysosomal vesicles in both immature and mature rat hippocampal neurons. Remarkably, the specialized microtubule environment of the axon initial segment (AIS) restricts KIF1A-driven transport of degradative, cathepsin D-positive lysosomal vesicles while permitting less degradative lysosomal vesicles to enter axons in mature neurons. Disrupting the AIS microtubule environment or expressing a hereditary spastic paraplegia type 30-associated KIF1A variant with increased affinity for GTP-tubulin bypasses this restriction, enabling degradative lysosomal vesicles to enter mature axons. In human iPSC-derived neurons, KIF1A depletion similarly disrupts axonal lysosomal vesicle transport, causes autophagic vesicle accumulation, and triggers axonal degeneration. Together, our findings reveal a developmentally regulated AIS gate that selectively controls KIF1A-driven lysosomal vesicle transport into axons and link its subversion to KIF1A-associated neurological disease.

## Introduction

Lysosomes are membrane-bound cytoplasmic organelles characterized by an acidic lumen containing a diverse array of hydrolytic enzymes that degrade various biomacromolecules, including proteins, lipids, nucleic acids, and carbohydrates.^1–3^ The lysosomal lumen is enclosed by a lipid bilayer enriched in specialized transmembrane proteins that maintain luminal acidity, mediate the export and recycling of degradation products, and regulate ionic and metabolic homeostasis. In addition, lysosomes act as key signaling hubs that integrate pathways controlling nutrient sensing, gene expression, and cellular growth.^1–3^

Although lysosomes are often regarded as a uniform population, they exhibit substantial heterogeneity in their morphology, composition, and function. To capture this heterogeneity, and unless otherwise specified, herein we refer collectively to these organelles as “lysosomal vesicles”. The diversity of lysosomal vesicles is particularly pronounced in mature neurons, where these organelles display distinct properties in different neuronal domains.^4,5^ In the soma, lysosomal vesicles generally exhibit the canonical features of degradative lysosomes, whereas those in the axon diverge from this profile in several respects. Most lysosomal vesicles undergoing anterograde axonal transport are relatively small, less acidic, and less degradative and contain variable amounts of non-lysosomal cargo, including synaptic vesicle proteins.^6–9^ Based on these properties, they have been proposed to represent biosynthetic carriers, lysosomal precursors, or immature lysosomes. In contrast, lysosomal vesicles undergoing retrograde axonal transport are generally larger, more acidic, and more degradative and are thought to represent mature lysosomes, late endosomes, autolysosomes, or amphisomes.^6,7,9–11^

The positioning and motility of lysosomes within different cell types^4,12^ are governed by their coupling to microtubule (MT) motors belonging to the kinesin and dynein families.^13,14^ In non- neuronal cells, both kinesin-1 (KIF5) and kinesin-3 (KIF1) family motors drive anterograde lysosome transport from the perinuclear region toward the cell periphery.^15–20^ In neurons, KIF5 has been shown to promote anterograde lysosomal vesicle transport into the axon.^21^ In contrast, the contribution of KIF1 motors in neuronal lysosomal vesicle transport remains poorly understood.

The *C. elegans* KIF1 motor UNC-104^22,23^ and the neuronally enriched mammalian KIF1A^24,25^ are the primary motors that drive the axonal anterograde transport of synaptic vesicle precursors (SVPs). A recent study demonstrated that inducible dimerization of exogenously expressed KIF1A tail and the KIF5C motor domains promotes the accumulation of lysosomal vesicles at axon terminals.^26^ However, whether full-length KIF1A itself mediates the axonal transport of lysosomal vesicles remains to be assessed. Defining the role of KIF1A in neuronal lysosomal transport is important not only for advancing our fundamental understanding of neuronal cell biology but also for elucidating the mechanisms underlying the broad spectrum of KIF1A- associated neurological disorders (KAND).^27–32^

To transport cargo within neurons, motors such as KIF1A must navigate a highly specialized and developmentally regulated MT cytoskeleton.^33,34^ In culture, embryonic rat hippocampal neurons undergo a well-characterized developmental program in which they progressively polarize and establish morphologically and functionally distinct axonal and dendritic compartments (Fig. 1a).^35,36^ These developmental transitions are accompanied by changes in MT organization and composition, as well as in tubulin post-translational modifications, that may differentially regulate motor-driven transport.^37–42^

**Figure 1.**
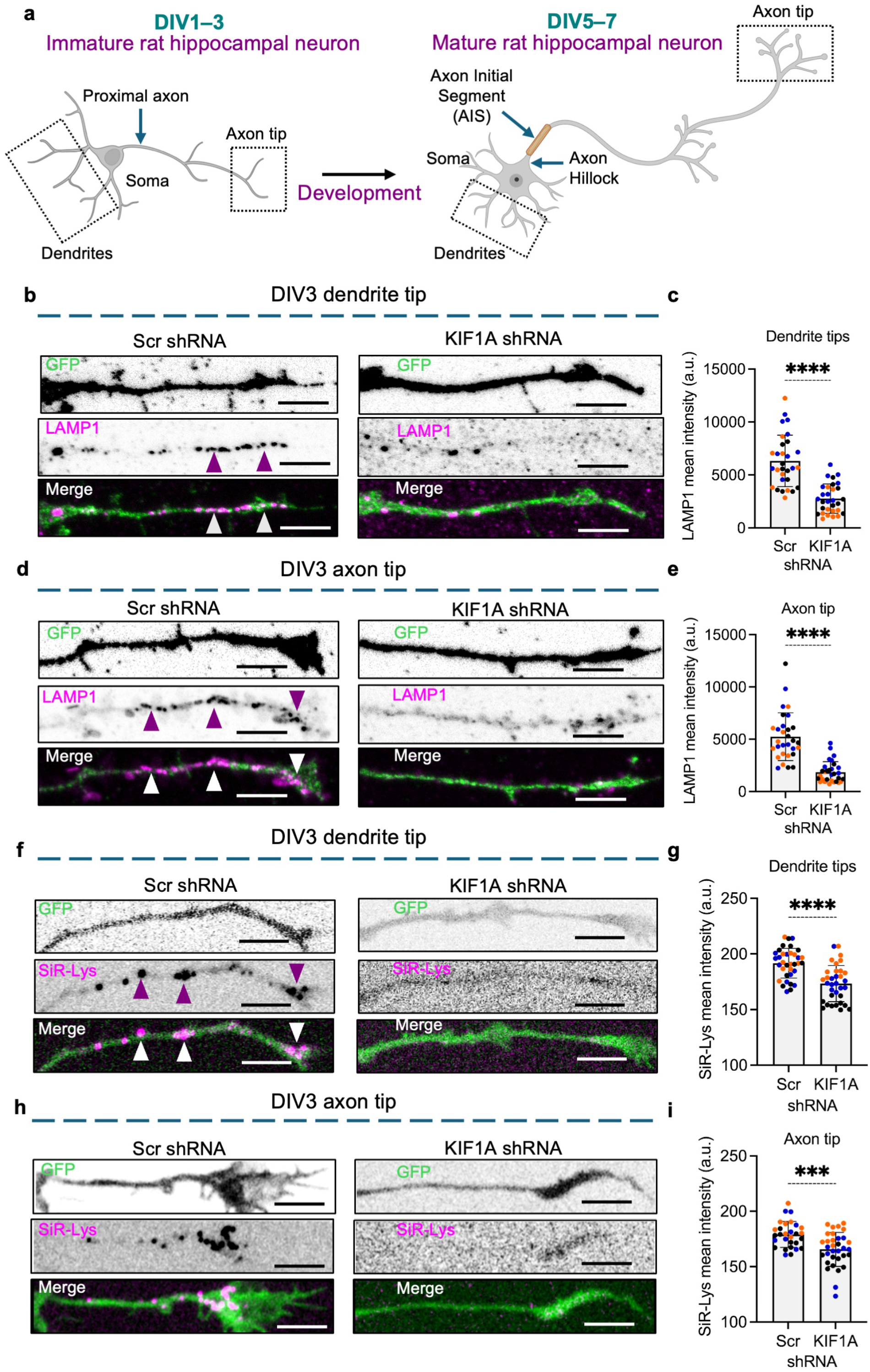
KIF1A is required for dendritic and axonal lysosome localization in immature rat hippocampal neurons. **a,** Schematic representation of immature (DIV1–3) and mature (DIV5–7) rat hippocampal neurons used in this study. Immature neurons are polarized with distinct dendritic and axonal compartments but lack a fully developed axon initial segment (AIS), whereas mature neurons contain a defined AIS adjacent to the axon hillock. Dendrite tips, the proximal axon/AIS, and axon tips analyzed throughout the study are indicated. **b,** Confocal fluorescence microscopy of dendrite tips from DIV3 rat hippocampal neurons transfected with plasmids encoding scrambled shRNA and GFP (Scr shRNA) or KIF1A shRNA and GFP, fixed, permeabilized, and immunostained for LAMP1. Single-channel images are shown in grayscale, and merged images in color with GFP in green and LAMP1 in magenta. Arrowheads indicate representative LAMP1-positive structures. Scale bars: 5 μm. **c,** Quantification showing that KIF1A shRNA significantly reduces LAMP1 intensity at dendrite tips. The graph shows individual data points and mean ± SD from 30 neurons per condition obtained from three independent cultures. Statistical significance was calculated by two-tailed Mann–Whitney test. ****P < 0.0001. **d,** Confocal fluorescence microscopy of axon tips from DIV3 rat hippocampal neurons expressing Scr shRNA or KIF1A shRNA and immunostained for LAMP1. Single-channel images are shown in grayscale, and merged images in color with GFP in green and LAMP1 in magenta. Arrowheads indicate representative LAMP1-positive structures. Scale bars: 5 μm. **e,** Quantification showing that KIF1A shRNA significantly reduces LAMP1 intensity at the axon tip. The graph shows individual data points and mean ± SD from 30 neurons per condition obtained from three independent cultures. Statistical significance was calculated by two-tailed Mann–Whitney test. ****P < 0.0001. **f,** Confocal live-cell imaging of dendrite tips from DIV3 rat hippocampal neurons expressing Scr shRNA or KIF1A shRNA and stained with the fluorogenic cathepsin D probe SiR-Lysosome (SiR-Lys). Single-channel images are shown in grayscale, and merged images in color with GFP in green and SiR-Lys in magenta. Arrowheads indicate representative SiR-Lys-positive structures. Scale bars: 5 μm. **g,** Quantification showing that KIF1A shRNA significantly reduces SiR-Lys intensity at dendrite tips. The graph shows individual data points and mean ± SD from 34–35 neurons per condition obtained from three independent cultures. Statistical significance was calculated by two-tailed Mann–Whitney test. ****P < 0.0001. **h,** Confocal live-cell imaging of axon tips from DIV3 rat hippocampal neurons expressing Scr shRNA or KIF1A shRNA and stained with SiR-Lys. Single-channel images are shown in grayscale, and merged images in color with GFP in green and SiR-Lys in magenta. Arrowheads indicate representative SiR-Lys-positive structures. Scale bars: 5 μm. **i,** Quantification showing that KIF1A shRNA significantly reduces SiR-Lys intensity at the axon tip. The graph shows individual data points and mean ± SD from 29–33 neurons per condition obtained from three independent cultures. Statistical significance was calculated by two-tailed Mann–Whitney test. ***P = 0.0006.

A defining feature of neuronal maturation is the formation of the axon initial segment (AIS), which emerges around day in vitro 3–4 (DIV3–4) in cultured rat hippocampal neurons.^43–45^ The AIS is a highly specialized structural and functional domain located in the proximal axon that serves as the primary site of action potential initiation. The scaffolding protein ankyrin-G (AnkG) is the master organizer of AIS architecture, anchoring membrane proteins such as neurofascin-186, voltage-gated sodium (Nav) and potassium (Kv) channels that are essential for neuronal excitability.^46–48^ AnkG also interacts with the underlying actin and MT cytoskeleton.^49^ AIS MTs are organized into fascicles of uniform polarity,^50–52^ with their plus ends oriented toward the distal axon.^37,53^ The AIS MTs contain tubulin in a conformation that resembles the GTP-bound state of the protein.^54^ They are also enriched in the end-tracking proteins EB1 and EB3, which associate with AnkG.^55,56^ Previous work has shown that KIF1A exhibits reduced affinity for GTP-tubulin-rich MTs and that the presence of EB3 on the MT lattice further weakens KIF1A–MT interactions.^57^ Given the distinctive MT lattice properties and enrichment of EB3 at the AIS, these findings raise the possibility that AIS formation during neuronal maturation creates a MT environment that regulates KIF1A function.

Here, we establish a role for KIF1A in lysosomal vesicle transport in rat hippocampal neurons and human iPSC-derived neurons. First, depletion analysis reveals that KIF1A is required for the dendritic and axonal localization of lysosomal vesicles in both immature and mature hippocampal neurons. Conversely, KIF1A overexpression enhances the accumulation of lysosomal vesicles in the dendrites and axons of immature neurons, as well as in the dendrites of mature hippocampal neurons. In the axon of mature neurons, however, KIF1A overexpression differentially affects the accumulation of two related lysosomal populations, increasing the abundance of lysosomal vesicles containing the lysosome-associated membrane protein 1 (LAMP1) without affecting that of lysosomal vesicles additionally containing the catalytically active protease, cathepsin D. This maturation-linked differential effect on lysosomal populations is due to the reduced ability of KIF1A to transport active cathepsin D-containing lysosomes through the AIS of mature neurons. Disrupting the MT environment of the AIS or enhancing KIF1A’s GTP-tubulin binding affinity via a mutation in the motor domain enhances KIF1A-driven transport of active cathepsin D containing lysosomes through the AIS, resulting in their increased abundance in the axons of mature neurons. Extending these findings to human neurons, we demonstrate that disruption of KIF1A in iPSC-derived i3Neurons impairs lysosomal vesicle transport in both immature and mature neurons, leading to accumulation of autophagic structures associated with mitochondria and eventual axonal degeneration. Our study thus establishes KIF1A as an essential motor for lysosomal vesicle transport in neurons and reveals how developmental changes in the MT environment at the AIS selectively regulate KIF1A-driven transport of degradative lysosomes into the axon of mature neurons.

## Results

### KIF1A is required for the dendritic and axonal localization of lysosomal vesicles in immature and mature hippocampal neurons

In this study, we investigated the role of KIF1A in the distribution and transport of two related populations of lysosomal vesicles in neurons: vesicles containing LAMP1 (lysosome-associated membrane protein 1), a marker of lysosomal vesicles irrespective of their degradative activity,^58^ and a more degradative subset additionally containing catalytically active cathepsin D.^59^ LAMP1-positive lysosomal vesicles were detected in fixed cells by immunofluorescence staining of endogenous LAMP1, whereas lysosomal vesicles containing catalytically active cathepsin D were detected in live cells using silicon-rhodamine-lysosome (SiR-Lys), a fluorogenic probe for active lysosomal cathepsin D.^60^

To examine whether KIF1A is required for lysosomal vesicle localization in immature neurons (Fig. 1a), we first transfected DIV1 rat hippocampal neurons with plasmids encoding either scrambled short hairpin RNA (Scr shRNA; control) or KIF1A-specific shRNA (KIF1A shRNA).^61^ Both plasmids also encoded green fluorescent protein (GFP) to identify transfected cells. We then analyzed the distribution of LAMP1 and SiR-Lys on DIV3. We observed that KIF1A depletion (KIF1A KD) (Fig. S1a) significantly reduced the amount of both LAMP1 (Fig. 1b-e, Fig. S1b) and SiR-Lys (Fig. 1f-i, Fig. S1c) signal at dendrite and axon tips of immature rat hippocampal neurons. Similarly, to investigate the role of KIF1A in lysosomal vesicle localization in mature hippocampal neurons (i.e., after AIS formation) (Fig. 1a), we transfected DIV5 rat hippocampal neurons with plasmids encoding Scr shRNA or KIF1A shRNA and analyzed the distribution of LAMP1 and SiR-Lys on DIV7. We observed that KIF1A KD significantly reduced the amount of both LAMP1 (Fig. 2a-d, Fig. S2a) and SiR-Lys (Fig. 2e-h, Fig. S2b) signal in the dendrites and axonal tips of mature rat hippocampal neurons.

**Figure 2.**
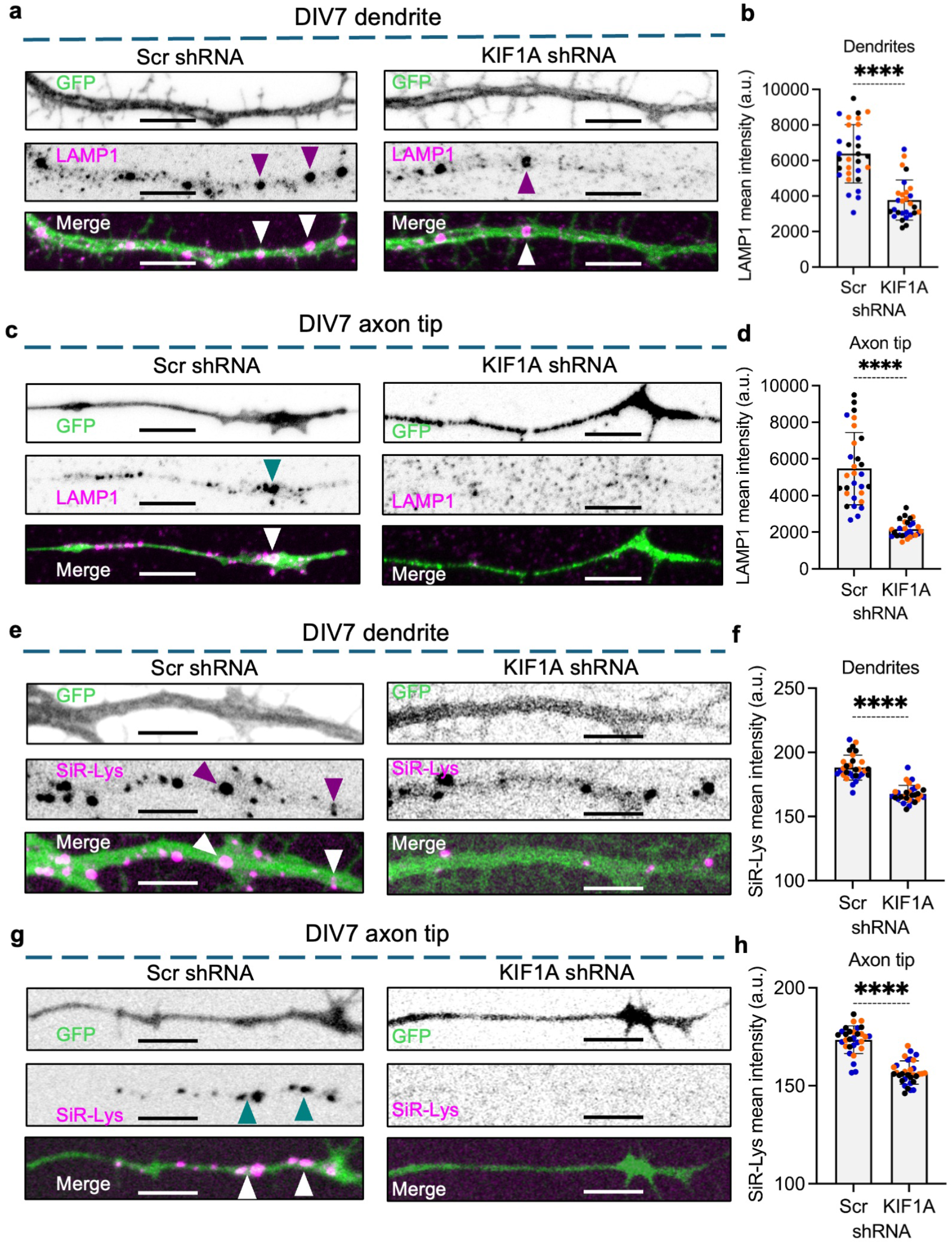
KIF1A is required for dendritic and axonal lysosome localization in mature rat hippocampal neurons. **a,** Confocal fluorescence microscopy of dendrites from DIV7 rat hippocampal neurons expressing Scr shRNA or KIF1A shRNA, fixed, permeabilized, and immunostained for LAMP1. Single-channel images are shown in grayscale, and merged images in color with GFP in green and LAMP1 in magenta. Arrowheads indicate representative LAMP1-positive structures. Scale bars: 5 μm. **b,** Quantification showing that KIF1A shRNA significantly reduces LAMP1 intensity in the dendrites. The graph shows individual data points and mean ± SD from 28–30 neurons per condition obtained from three independent cultures. Statistical significance was calculated by two-tailed Mann–Whitney test. ****P < 0.0001. **c,** Confocal fluorescence microscopy of axon tips from DIV7 rat hippocampal neurons expressing Scr shRNA or KIF1A shRNA and immunostained for LAMP1. Single-channel images are shown in grayscale, and merged images in color with GFP in green and LAMP1 in magenta. Arrowheads indicate representative LAMP1-positive structures. Scale bars: 5 μm. **d,** Quantification showing that KIF1A shRNA significantly reduces LAMP1 intensity at the axon tip. The graph shows individual data points and mean ± SD from 28–30 neurons per condition obtained from three independent cultures. Statistical significance was calculated by two-tailed Mann–Whitney test. ****P < 0.0001. **e,** Confocal live-cell imaging of dendrites from DIV7 rat hippocampal neurons expressing Scr shRNA or KIF1A shRNA and stained with SiR-Lys. Single-channel images are shown in grayscale, and merged images in color with GFP in green and SiR-Lys in magenta. Arrowheads indicate representative SiR-Lys-positive structures. Scale bars: 5 μm. **f,** Quantification showing that KIF1A shRNA significantly reduces SiR-Lys intensity in the dendrites. The graph shows individual data points and mean ± SD from 30 neurons per condition obtained from three independent cultures. Statistical significance was calculated by two-tailed Mann–Whitney test. ****P < 0.0001. **g,** Confocal live-cell imaging of axon tips from DIV7 rat hippocampal neurons expressing Scr shRNA or KIF1A shRNA and stained with SiR-Lys. Single-channel images are shown in grayscale, and merged images in color with GFP in green and SiR-Lys in magenta. Arrowheads indicate representative SiR-Lys-positive structures. Scale bars: 5 μm. **h,** Quantification showing that KIF1A shRNA significantly reduces SiR-Lys intensity at the axon tip. The graph shows individual data points and mean ± SD from 31–32 neurons per condition obtained from three independent cultures. Statistical significance was calculated by two-tailed Mann–Whitney test. ****P < 0.0001.

These results indicate that KIF1A is an essential motor for the distribution of lysosomal vesicles in the dendritic and axonal compartments of both immature and mature hippocampal neurons.

### KIF1A overexpression has differential effects on lysosomal vesicle localization in immature and mature hippocampal neurons

We next investigated the ability of KIF1A overexpression to enhance the accumulation of lysosomal vesicles in the dendrites and axons of immature hippocampal neurons. We first transfected DIV2 rat hippocampal neurons with plasmids encoding either GFP (control) or GFP- tagged mouse KIF1A (GFP-KIF1A).^20^ On DIV3, we fixed the neurons, immunostained them for endogenous LAMP1, and analyzed the distributions of GFP/GFP-KIF1A and LAMP1 by confocal fluorescence microscopy. We observed that GFP-KIF1A colocalized with LAMP1 in the somatodendritic compartment and increased the accumulation of LAMP1 at dendritic tips (Fig. 3a,b; Fig. S3a) and axonal tips (Fig. 3c,d). Similarly, we transfected DIV2 rat hippocampal neurons with plasmids encoding mNeonGreen (mNG) or mNeonGreen-tagged KIF1A (mNG- KIF1A) and observed that, at DIV3, mNG-KIF1A colocalized with SiR-Lys and increased the accumulation of SiR-Lys at both dendritic tips (Fig. 3e,f, Fig. S3b) and axonal tips (Fig. 3g,h). These results establish that, in immature neurons, KIF1A can drive the transport of lysosomal vesicles toward MT plus ends, which are uniformly oriented toward the distal ends of both dendrites^38^ and axons.^37,53^

**Figure 3.**
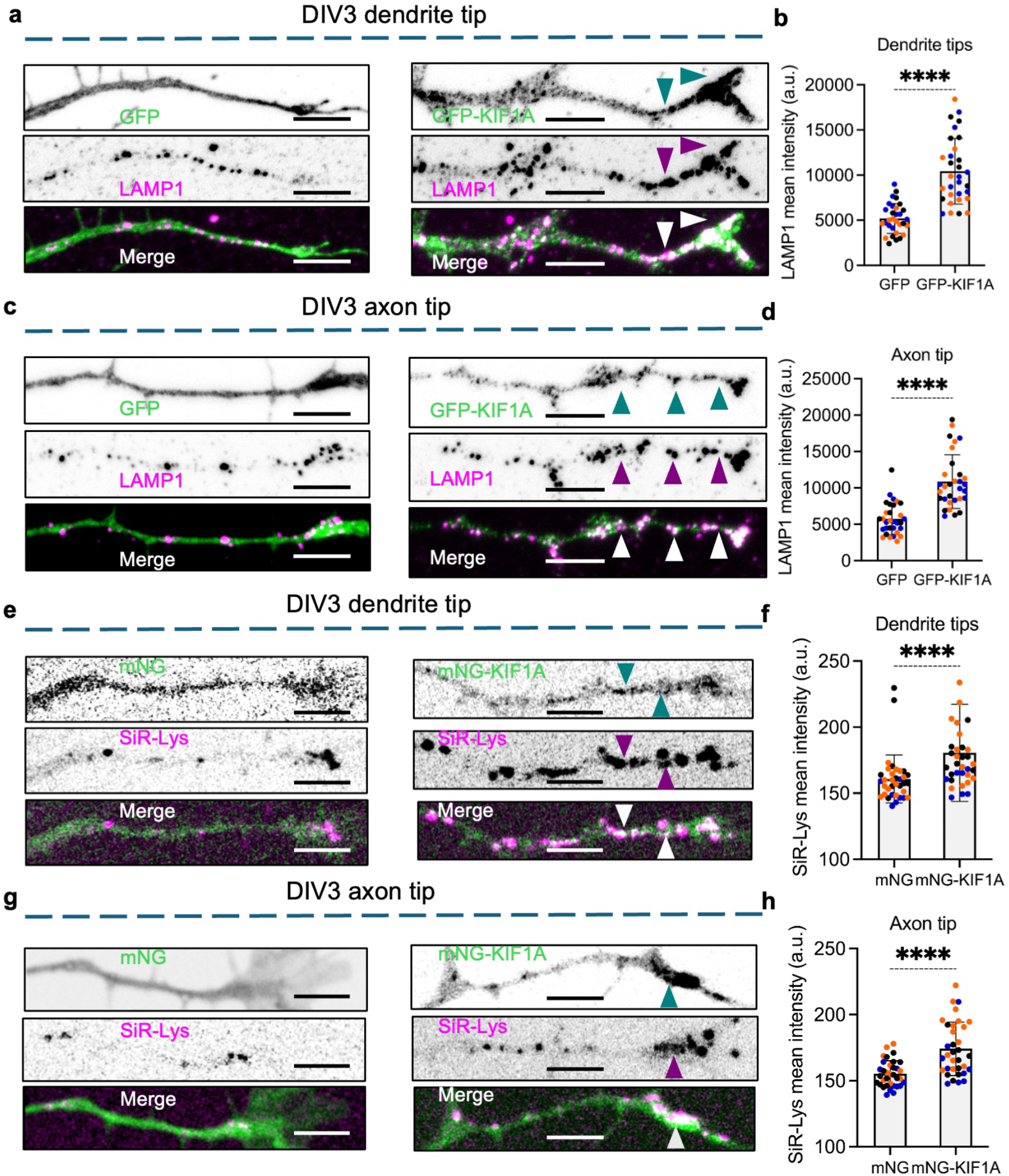
KIF1A expression increases lysosome accumulation in the dendrites and axons of immature rat hippocampal neurons. **a,** Confocal fluorescence microscopy of dendrite tips from DIV3 rat hippocampal neurons expressing GFP or GFP-KIF1A, fixed, permeabilized, and immunostained for LAMP1. Single- channel images are shown in grayscale, and merged images in color with GFP in green and LAMP1 in magenta. Arrowheads indicate representative GFP-KIF1A- and LAMP1-positive structures. Scale bars: 5 μm. **b,** Quantification showing that GFP-KIF1A expression significantly increases LAMP1 intensity at dendrite tips. The graph shows individual data points and mean ± SD from 31 neurons per condition obtained from three independent cultures. Statistical significance was calculated by unpaired Welch’s t-test. ****P < 0.0001. **c,** Confocal fluorescence microscopy of axon tips from DIV3 rat hippocampal neurons expressing GFP or GFP-KIF1A and immunostained for LAMP1. Single-channel images are shown in grayscale, and merged images in color with GFP in green and LAMP1 in magenta. Arrowheads indicate representative GFP-KIF1A- and LAMP1-positive structures. Scale bars: 5 μm. **d,** Quantification showing that GFP-KIF1A expression significantly increases LAMP1 intensity at the axon tip. The graph shows individual data points and mean ± SD from 31 neurons per condition obtained from three independent cultures. Statistical significance was calculated by two-tailed Mann–Whitney test. ****P < 0.0001. **e,** Confocal live-cell imaging of dendrite tips from DIV3 rat hippocampal neurons expressing mNeonGreen (mNG) or mNeonGreen-KIF1A (mNG-KIF1A) and stained with SiR-Lys. Single- channel images are shown in grayscale, and merged images in color with mNG or mNG-KIF1A in green and SiR-Lys in magenta. Arrowheads indicate representative mNG-KIF1A- and SiR- Lys-positive structures. Scale bars: 5 μm. **f,** Quantification showing that mNG-KIF1A expression significantly increases SiR-Lys intensity at dendrite tips. The graph shows individual data points and mean ± SD from 34–35 neurons per condition obtained from three independent cultures. Statistical significance was calculated by two-tailed Mann–Whitney test. ****P < 0.0001. **g,** Confocal live-cell imaging of axon tips from DIV3 rat hippocampal neurons expressing mNG or mNG-KIF1A and stained with SiR-Lys. Single-channel images are shown in grayscale, and merged images in color with mNG or mNG-KIF1A in green and SiR-Lys in magenta. Arrowheads indicate representative mNG-KIF1A- and SiR-Lys-positive structures. Scale bars: 5 μm. **h,** Quantification showing that mNG-KIF1A expression significantly increases SiR-Lys intensity at axon tips. The graph shows individual data points and mean ± SD from 34 neurons per condition obtained from three independent cultures. Statistical significance was calculated by two-tailed Mann–Whitney test. ****P < 0.0001.

Next, we examined whether KIF1A overexpression drives the transport of lysosomal vesicles in mature (DIV5–7) rat hippocampal neurons, which are characterized by the presence of a well- defined axon initial segment (AIS) and greater dendritic and axonal complexity (Fig. 1a). To this end, we transfected DIV6 rat hippocampal neurons with plasmids encoding GFP or GFP- KIF1A, fixed the neurons on DIV7, immunostained them for endogenous LAMP1 and AnkG, and analyzed the distributions of GFP/GFP-KIF1A and LAMP1 by confocal fluorescence microscopy. In GFP-expressing cells, LAMP1 puncta were uniformly distributed along the dendrites. In contrast, in GFP-KIF1A-expressing cells, GFP-KIF1A and LAMP1 accumulated in fewer, larger clusters along the dendrites and at dendritic tips (Fig. 4a,b; Fig. S4a). This distribution is consistent with the mixed polarity of MTs in mature dendrites and the accumulation of LAMP1-positive vesicles at MT plus ends along the dendrites.^37^ Moreover, GFP-KIF1A overexpression significantly increased the accumulation of LAMP1-positive lysosomal vesicles at the axon tip (Fig. 4c,d), similar to the effect observed in immature neurons (Fig. 3c,d).

**Figure 4.**
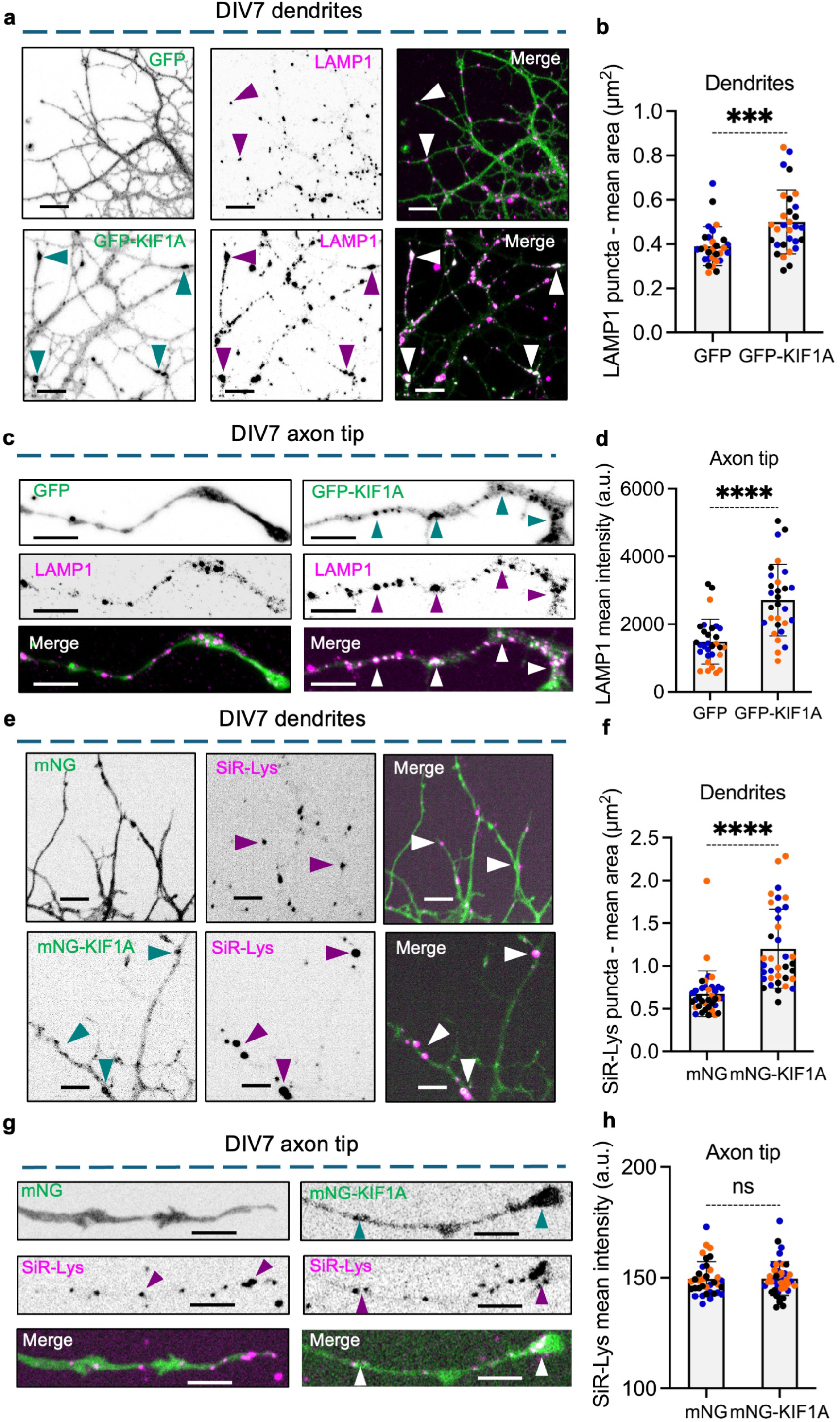
KIF1A expression differentially affects lysosome populations in mature rat hippocampal neurons. **a,** Confocal fluorescence microscopy of the dendrites of DIV7 rat hippocampal neurons expressing GFP or GFP-KIF1A, fixed, permeabilized, and immunostained for LAMP1. Single- channel images are shown in grayscale, and merged images in color with GFP in green and LAMP1 in magenta. Arrowheads indicate representative LAMP1-positive structures. Scale bars: 20 μm. **b,** Quantification showing that GFP-KIF1A expression significantly increases the mean area of LAMP1-positive puncta within dendrites. The graph shows individual data points and mean ± SD from 29–30 neurons per condition obtained from three independent cultures. Statistical significance was calculated by two-tailed Mann–Whitney test. ***P = 0.0004. **c,** Confocal fluorescence microscopy of axon tips from DIV7 rat hippocampal neurons expressing GFP or GFP-KIF1A and immunostained for LAMP1. Single-channel images are shown in grayscale, and merged images in color with GFP in green and LAMP1 in magenta. Arrowheads indicate representative GFP-KIF1A- and LAMP1-positive structures. Scale bars: 5 μm. **d,** Quantification showing that GFP-KIF1A expression significantly increases LAMP1 intensity at axon tips. The graph shows individual data points and mean ± SD from 29–30 neurons per condition obtained from three independent cultures. Statistical significance was calculated by two-tailed Mann–Whitney test. ****P < 0.0001. **e,** Confocal live-cell imaging of the dendrites of DIV7 rat hippocampal neurons expressing mNG or mNG-KIF1A and stained with SiR-Lys. Single-channel images are shown in grayscale, and merged images in color with mNG or mNG-KIF1A in green and SiR-Lys in magenta. Arrowheads indicate representative mNG-KIF1A- and SiR-Lys-positive structures. Scale bars: 20 μm. **f,** Quantification showing that mNG-KIF1A expression significantly increases the area of SiR- Lys-positive puncta within dendrites. The graph shows individual data points and mean ± SD from 36–37 neurons per condition obtained from three independent cultures. Statistical significance was calculated by two-tailed Mann–Whitney test. ****P < 0.0001. **g,** Confocal live-cell imaging of axon tips from DIV7 rat hippocampal neurons expressing mNG or mNG-KIF1A and stained with SiR-Lys. Single-channel images are shown in grayscale, and merged images in color with mNG or mNG-KIF1A in green and SiR-Lys in magenta. Scale bars: 5 μm. **h,** Quantification showing that mNG-KIF1A expression does not significantly increase SiR-Lys intensity at axon tips. The graph shows individual data points and mean ± SD from 35–42 neurons per condition obtained from three independent cultures. Statistical significance was calculated by two-tailed Mann–Whitney test. ns, not significant.

We then asked whether KIF1A similarly drives the transport of lysosomes containing active cathepsin D in mature hippocampal neurons. We transfected DIV6 neurons with plasmids encoding mNG (control) or mNG-KIF1A and, on DIV7, stained them with SiR-Lys and a CF 555-conjugated extracellular anti-neurofascin antibody (NF-555) to label the AIS for live-cell microscopy. We observed that, in DIV7 neurons, mNG-KIF1A and SiR-Lys colocalized in large puncta along the dendrites (Fig. 4e,f). Surprisingly, although mNG-KIF1A fluorescence was readily detected at axon tips, mNG-KIF1A overexpression did not significantly increase the accumulation of SiR-Lys at axon tips in mature hippocampal neurons (Fig. 4g,h).

We next tested whether this effect was specific to SiR-Lys-positive lysosomal vesicles. We compared the accumulation of the synaptic vesicle protein synaptophysin (SYP) at axon tips of DIV7 neurons overexpressing GFP or GFP-KIF1A and found that GFP-KIF1A overexpression significantly increased SYP accumulation (Fig. S4c,d).

These results establish that, in mature neurons, KIF1A overexpression differentially affects the axonal localization of lysosomal vesicle populations with distinct degradative activity while retaining its ability to transport SVPs into the axon. We hypothesized that this differential effect might result from preferential transport of less degradative, LAMP1-positive but SiR-Lys- negative lysosomal vesicles into the axons of mature hippocampal neurons.

### KIF1A preferentially moves less degradative lysosomal vesicles into the axons of mature neurons

To test our hypothesis, we compared the effect of KIF1A expression on the movement of SiR- Lys-positive vesicles through the proximal axon of immature neurons with their movement through the AIS of mature neurons. We generated kymographs of the proximal axon from dual-color spinning-disk confocal microscopy videos of DIV3 hippocampal neurons expressing mNG or mNG-KIF1A and labeled with SiR-Lys. We observed mNG-KIF1A tracks moving through the proximal axon of DIV3 neurons, most of which colocalized with SiR-Lys tracks (Fig. 5a).

**Figure 5.**
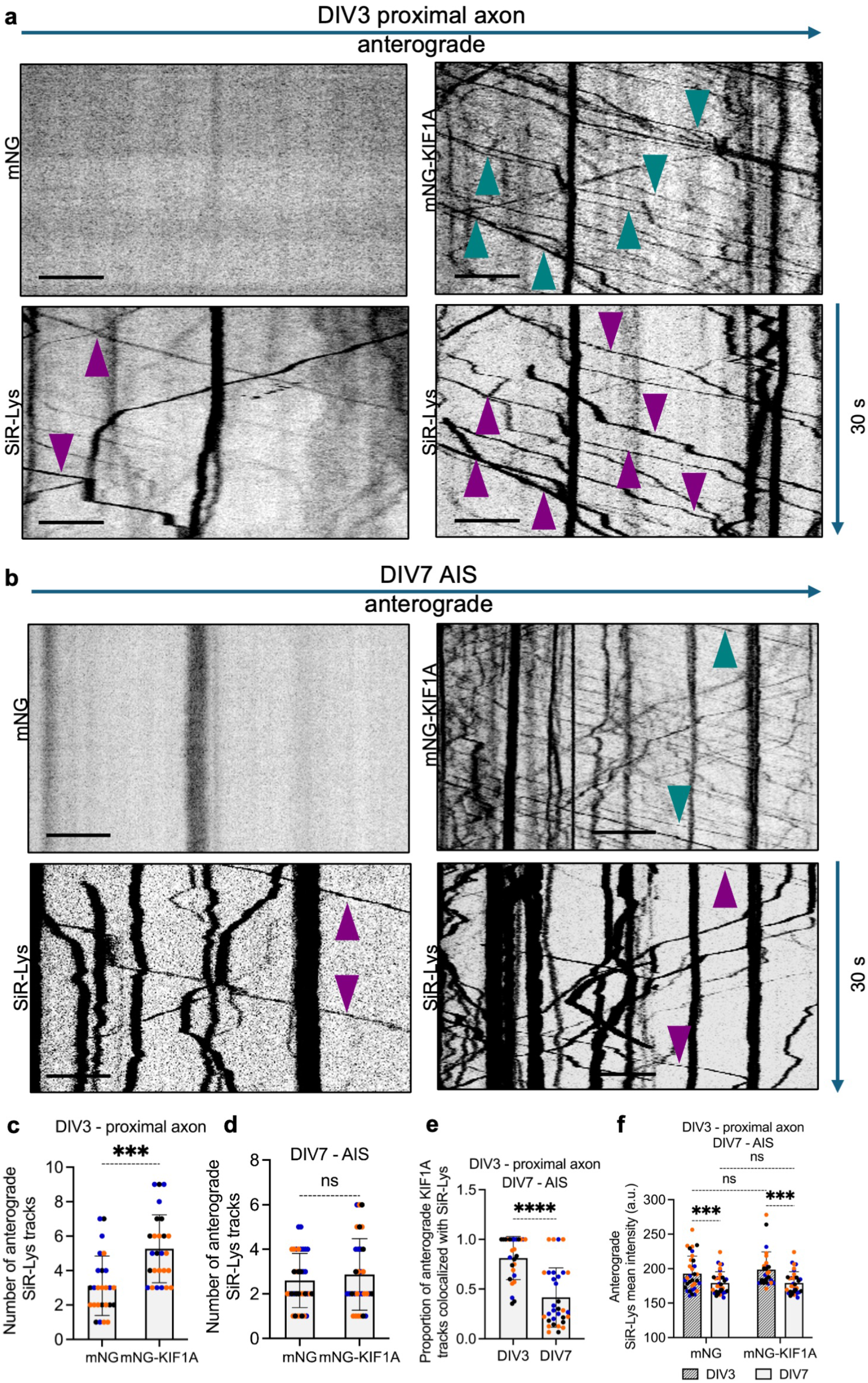
Neuronal maturation restricts KIF1A-driven transport of degradative lysosomes into the axon. **a,** Kymographs generated from live videos of the proximal axon of DIV3 rat hippocampal neurons expressing mNG or mNG-KIF1A and stained with SiR-Lys. Teal arrowheads indicate representative mNG-KIF1A-positive anterograde tracks, and magenta arrowheads indicate SiR- Lys-positive anterograde tracks. Scale bars: 5 μm. **b,** Kymographs generated from live videos of the AIS of DIV7 rat hippocampal neurons expressing mNG or mNG-KIF1A and stained with SiR-Lys. NF-555 staining was used to identify the AIS. Teal arrowheads indicate representative mNG-KIF1A-positive anterograde tracks, and magenta arrowheads indicate SiR-Lys-positive anterograde tracks. Scale bars: 5 μm. **c,** Quantification showing that mNG-KIF1A expression significantly increases the number of anterograde SiR-Lys tracks through the proximal axon of DIV3 neurons. The graph shows individual data points and mean ± SD from 26 neurons per condition obtained from three independent cultures. Statistical significance was calculated by two-tailed Welch’s t-test. ***P < 0.001. **d,** Quantification showing that mNG-KIF1A expression does not significantly increase the number of anterograde SiR-Lys tracks through the AIS of DIV7 neurons. The graph shows individual data points and mean ± SD from 31–35 neurons per condition obtained from three independent cultures. Statistical significance was calculated by two-tailed Welch’s t-test. ns, P = 0.4640. **e,** Quantification showing that the proportion of anterograde mNG-KIF1A tracks colocalizing with SiR-Lys is significantly lower at the AIS of DIV7 neurons than in the proximal axon of DIV3 neurons. The graph shows individual data points and mean ± SD from 27–32 neurons obtained from three independent cultures. Statistical significance was calculated by two-tailed Mann–Whitney test. ****P < 0.0001. **f,** Quantification of the mean intensity of anterograde SiR-Lys tracks through the proximal axon of DIV3 neurons and the AIS of DIV7 neurons expressing mNG or mNG-KIF1A. Anterograde SiR-Lys tracks are less intense in mature DIV7 neurons than in immature DIV3 neurons. The graph shows individual data points and mean ± SD from 27–35 neurons obtained from three independent cultures. Statistical significance was calculated by Kruskal–Wallis test followed by Dunn’s multiple-comparisons test. ****P < 0.0001; ***P < 0.001; ns, not significant.

Furthermore, mNG-KIF1A expression significantly increased the number of anterograde SiR- Lys tracks moving through the proximal axon of DIV3 neurons (Fig. 5a,c; Video-S1). Similarly, we generated kymographs of the AIS, identified by Neurofascin-555 staining, from dual-color spinning-disk confocal microscopy videos of DIV7 hippocampal neurons expressing mNG or mNG-KIF1A and stained with SiR-Lys. In mature neurons, we also observed mNG-KIF1A tracks moving through the AIS, but very few colocalized with SiR-Lys tracks (Fig. 5b).

Moreover, in contrast to DIV3 neurons, mNG-KIF1A expression did not significantly increase the number of anterograde SiR-Lys tracks moving through the AIS of DIV7 neurons (Fig. 5b,d; Video-S2). The proportion of anterograde KIF1A tracks colocalized with SiR-Lys was also significantly lower in the AIS of DIV7 neurons than in the proximal axon of DIV3 neurons (Fig. 5e), and the intensity of anterogradely moving SiR-Lys tracks was significantly lower in mature neurons (Fig. 5f).

We then analyzed differences in the movement of lysosomal vesicles into the axons of immature and mature neurons under endogenous levels of KIF1A expression. We transfected DIV2 or DIV6 neurons with a plasmid encoding mNeonGreen-tagged LAMP1 (LAMP1-mNG), stained them with SiR-Lys on DIV3 or DIV7, respectively, and performed dual-color spinning-disk confocal microscopy of the proximal axon/AIS region (Fig. S5a) and generated kymographs of the proximal axon in DIV3 neurons and the AIS in DIV7 neurons (Fig. S5b). Kymograph analysis revealed that the proportion of anterograde LAMP1 tracks colocalized with SiR-Lys was significantly lower in the AIS of DIV7 neurons than in the proximal axon of DIV3 neurons (Fig. S5b,c; Video-S3). In contrast, the proportion of SiR-Lys tracks colocalized with LAMP1- mNG was close to one and did not differ significantly between DIV3 and DIV7 neurons (Fig. S5d), consistent with SiR-Lys-positive vesicles representing a subset of LAMP1-positive lysosomal vesicles. Importantly, the intensity of anterogradely moving SiR-Lys tracks was significantly lower in mature neurons (Fig. S5e), similar to our observations of SiR-Lys movement upon KIF1A overexpression. Together, these results suggest that AIS formation developmentally regulates KIF1A-driven lysosomal transport, favoring the delivery of less degradative lysosomal vesicles into the axons of mature neurons.

### Ankyrin-G-mediated EB protein localization at the AIS regulates KIF1A-driven transport of degradative lysosomes

The MT lattice at the AIS is enriched in the plus-end-binding proteins EB1 and EB3.^55,56,62^ Moreover, EB3 reduces the MT-binding affinity of KIF1A.^57^ We therefore hypothesized that enrichment of EB proteins at the mature AIS restricts KIF1A-dependent transport of SiR-Lys- positive lysosomal vesicles into the axon. We first investigated the interdependence of EB protein and AnkG localization at the AIS of DIV5–7 hippocampal neurons. AnkG depletion between DIV5 and DIV7 led to a marked reduction in EB1 enrichment at the AIS, whereas combined depletion of EB1 and EB3 markedly reduced EB1 staining while discernible AnkG staining remained at the AIS (Fig. S6). Consistent with these observations, AnkG depletion caused a marked loss of NF-555 staining at the AIS, whereas neurons depleted of EB1+EB3 retained NF-555 staining (Fig. 6a), indicating that EB1+EB3 depletion did not completely disrupt AIS organization under these conditions.^55^

**Figure 6.**
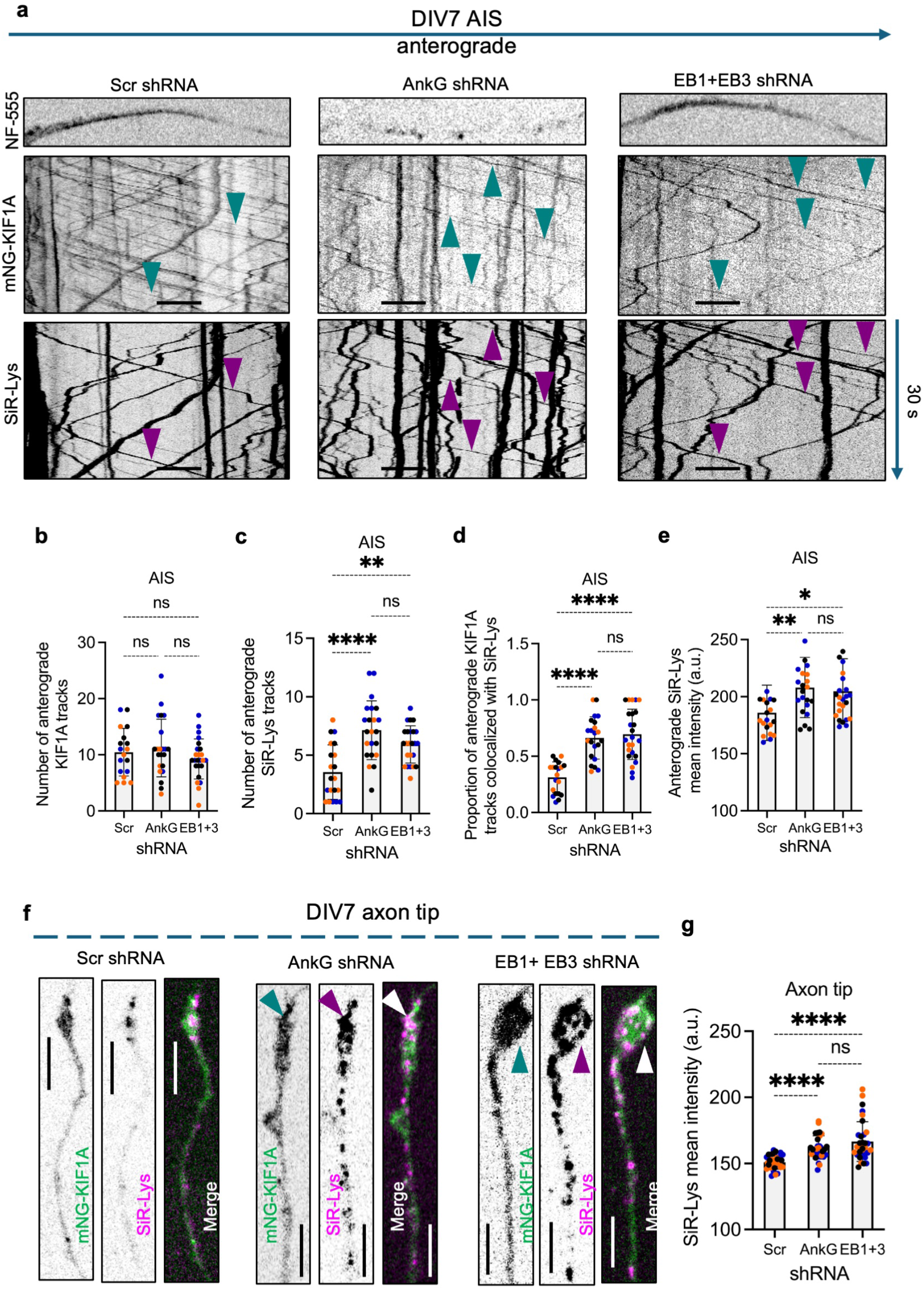
Ankyrin-G-mediated EB protein localization at the AIS restricts KIF1A-driven transport of degradative lysosomes. **a,** Kymographs generated from live videos of the AIS of DIV7 rat hippocampal neurons expressing Scr shRNA, ankyrin-G (AnkG) shRNA, or EB1+EB3 shRNA together with mNG- KIF1A-WT and stained with SiR-Lys. Neurofascin-555 (NF-555) was used to identify the AIS. Teal arrowheads indicate representative mNG-KIF1A-WT-positive anterograde tracks, and magenta arrowheads indicate SiR-Lys-positive anterograde tracks. Scale bars: 5 μm. **b,** Quantification showing that expression of AnkG shRNA or EB1+EB3 shRNA does not significantly alter the number of anterograde mNG-KIF1A-WT tracks through the AIS. The graph shows individual data points and mean ± SD from 20–24 neurons per condition obtained from three independent cultures. Statistical significance was calculated by ordinary one-way ANOVA followed by Tukey’s multiple-comparisons test. ns, not significant. **c,** Quantification showing that expression of AnkG shRNA or EB1+EB3 shRNA significantly increases the number of anterograde SiR-Lys tracks through the AIS compared with Scr shRNA. The graph shows individual data points and mean ± SD from 20–24 neurons per condition obtained from three independent cultures. Statistical significance was calculated by ordinary one-way ANOVA followed by Tukey’s multiple-comparisons test. **P < 0.005; ****P < 0.0001; ns, not significant. **d,** Quantification showing that the proportion of anterograde mNG-KIF1A-WT tracks colocalizing with SiR-Lys is significantly increased following expression of AnkG shRNA or EB1+EB3 shRNA compared with Scr shRNA. The graph shows individual data points and mean ± SD from 20–24 neurons per condition obtained from three independent cultures. Statistical significance was calculated by ordinary one-way ANOVA followed by Tukey’s multiple-comparisons test. ****P < 0.0001; ns, not significant. **e,** Quantification of the mean intensity of anterograde SiR-Lys tracks through the AIS in neurons expressing Scr shRNA, AnkG shRNA, or EB1+EB3 shRNA. SiR-Lys tracks are brighter following AnkG or EB1+EB3 depletion. The graph shows individual data points and mean ± SD from 20–24 neurons per condition obtained from three independent cultures. Statistical significance was calculated by Kruskal–Wallis test followed by Dunn’s multiple-comparisons test. **P < 0.01; *P < 0.05; ns, not significant. **f,** Confocal live-cell imaging of axon tips from DIV7 rat hippocampal neurons expressing Scr shRNA, AnkG shRNA, or EB1+EB3 shRNA together with mNG-KIF1A-WT and stained with SiR-Lys. Single-channel images are shown in grayscale, and merged images in color with mNG- KIF1A-WT in green and SiR-Lys in magenta. Arrowheads indicate representative mNG-KIF1A- WT- and SiR-Lys-positive structures. Scale bars: 5 μm. **g,** Quantification showing that expression of AnkG shRNA or EB1+EB3 shRNA significantly increases SiR-Lys intensity at axon tips compared with Scr shRNA. The graph shows individual data points and mean ± SD from 30–32 neurons per condition obtained from three independent cultures. Statistical significance was calculated by ordinary one-way ANOVA followed by Tukey’s multiple-comparisons test. ****P < 0.0001; ns, not significant.

To determine how these perturbations affect KIF1A-driven transport of degradative lysosomes through the AIS, we co-transfected DIV5 hippocampal neurons with plasmids encoding scrambled shRNA, AnkG shRNA, or a combination of EB1- and EB3-specific shRNAs (EB1+EB3 shRNA), together with a plasmid encoding mNG-KIF1A. On DIV7, we generated kymographs of mNG-KIF1A and SiR-Lys movement through the AIS (Fig. 6a; Video-S4). Depletion of AnkG or EB1+EB3 did not significantly affect the number of anterograde KIF1A tracks through the AIS (Fig. 6b) but significantly increased the number of anterograde SiR-Lys tracks (Fig. 6c). Moreover, depletion of AnkG or EB1+EB3 significantly increased both the proportion of anterograde KIF1A tracks that colocalized with SiR-Lys (Fig. 6d) and the average intensity of anterogradely moving SiR-Lys tracks (Fig. 6e). Finally, depletion of AnkG or EB1+EB3 significantly increased the accumulation of SiR-Lys at the axon tip (Fig. 6f,g), consistent with increased KIF1A-driven transport of more degradative lysosomes from the soma into the axon. Together, these observations indicate that disrupting EB protein enrichment at the AIS permits KIF1A to transport more SiR-Lys-positive lysosomal vesicles into the mature axon without increasing overall anterograde KIF1A traffic.

### KIF1A-T258M increases the transport of degradative lysosomes into mature axons

We then asked whether increasing KIF1A interaction with the GTP-tubulin-enriched MT lattice at the AIS^54^ could similarly promote the transport of degradative lysosomes through the AIS. To test this, we used the disease-associated KIF1A-T258M mutant,^63^ which has increased affinity for GTP-tubulin.^57^ We transfected DIV6 hippocampal neurons with plasmids encoding mNG, mNG-KIF1A-WT, or mNG-KIF1A-T258M, labeled the cells with SiR-Lys on DIV7, and performed dual-color spinning-disk confocal microscopy.

Kymograph analysis of the AIS revealed that KIF1A-T258M overexpression did not significantly alter the number of anterograde KIF1A tracks relative to KIF1A-WT overexpression (Fig. 7a,b). However, KIF1A-T258M overexpression significantly increased the number of anterograde SiR- Lys tracks relative to both mNG and KIF1A-WT (Fig. 7a,c). Moreover, a significantly greater proportion of anterograde KIF1A-T258M tracks colocalized with SiR-Lys than KIF1A-WT tracks (Fig. 7d; Video-S5). Anterogradely moving SiR-Lys tracks were also significantly brighter in KIF1A-T258M-expressing neurons than in KIF1A-WT- or mNG-expressing neurons (Fig. 7e). Interestingly, despite transporting more SiR-Lys-positive vesicles through the AIS, KIF1A- T258M moved significantly slower than KIF1A-WT within the AIS (Fig. 7f), as reported in previous *in vitro* studies.^57^ Consistent with the increased transport of SiR-Lys-positive vesicles through the AIS, expression of KIF1A-T258M also significantly increased SiR-Lys accumulation at the axon tips of DIV7 neurons, whereas KIF1A-WT did not significantly increase axon-tip SiR-Lys accumulation compared with mNG (Fig. 7g,h). Together, these results indicate that increased GTP-tubulin affinity enables KIF1A to transport more degradative, SiR-Lys-positive lysosomal vesicles through the AIS and into mature axons, without increasing the overall number of anterograde KIF1A tracks. Our data thus support a model in which the MT environment of the AIS modulates KIF1A-dependent lysosomal cargo transport into the axon.

**Figure 7.**
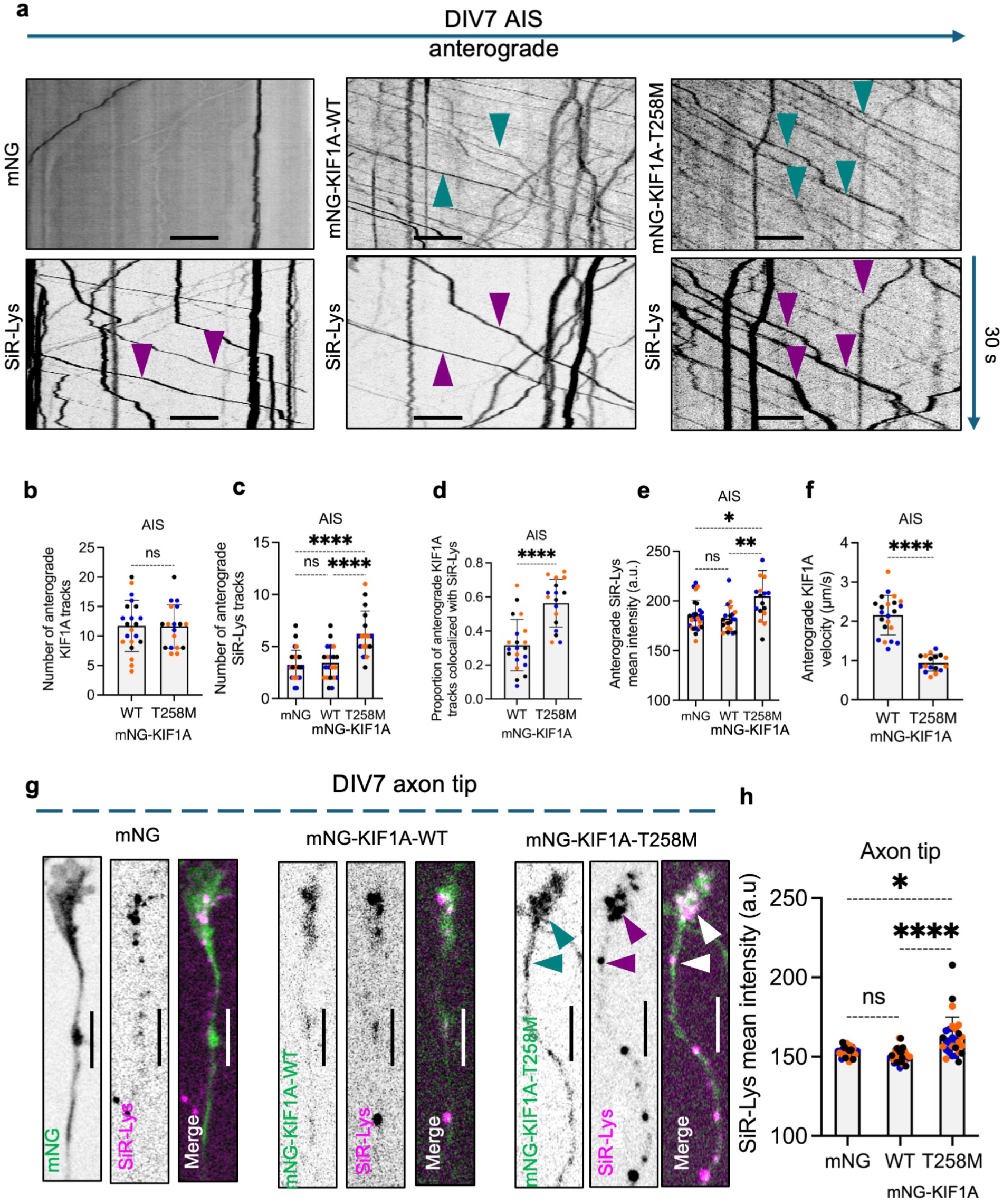
KIF1A-T258M increases the anterograde transport of degradative lysosomes through the AIS. **a,** Kymographs generated from live videos of the AIS of DIV7 rat hippocampal neurons expressing mNG, mNG-KIF1A-WT, or mNG-KIF1A-T258M and stained with SiR-Lys. Teal arrowheads indicate representative mNG-KIF1A-positive anterograde tracks, and magenta arrowheads indicate SiR-Lys-positive anterograde tracks. Scale bars: 5 μm. **b,** Quantification showing that the number of anterograde KIF1A tracks through the AIS does not significantly differ between neurons expressing mNG-KIF1A-WT and mNG-KIF1A-T258M. The graph shows individual data points and mean ± SD from 17–21 neurons obtained from three independent cultures. Statistical significance was calculated by two-tailed unpaired Welch’s t-test. ns, not significant. **c,** Quantification showing that expression of mNG-KIF1A-T258M significantly increases the number of anterograde SiR-Lys tracks through the AIS compared with mNG or mNG-KIF1A- WT. The graph shows individual data points and mean ± SD from 17–27 neurons per condition obtained from three independent cultures. Statistical significance was calculated by ordinary one-way ANOVA followed by Tukey’s multiple-comparisons test. ****P < 0.0001; ns, not significant. **d,** Quantification showing that the proportion of anterograde KIF1A tracks colocalizing with SiR-Lys is significantly increased for mNG-KIF1A-T258M compared with mNG-KIF1A-WT. The graph shows individual data points and mean ± SD from 17–21 neurons per condition obtained from three independent cultures. Statistical significance was calculated by two-tailed unpaired Welch’s t-test. ****P < 0.0001. **e,** Quantification of the mean intensity of anterograde SiR-Lys tracks through the AIS in neurons expressing mNG, mNG-KIF1A-WT, or mNG-KIF1A-T258M. SiR-Lys tracks are brighter in mNG-KIF1A-T258M-expressing neurons. The graph shows individual data points and mean ± SD from 17–21 neurons per condition obtained from three independent cultures. Statistical significance was calculated by Kruskal–Wallis test followed by Dunn’s multiple-comparisons test. **P < 0.01; *P < 0.05; ns, not significant. **f,** Quantification showing that the mean velocity of anterograde mNG-KIF1A-T258M tracks through the AIS is significantly reduced compared with mNG-KIF1A-WT tracks. The graph shows individual data points and mean ± SD from 17–22 neurons per condition obtained from three independent cultures. Statistical significance was calculated by two-tailed unpaired Welch’s t-test. ****P < 0.0001. **g,** Confocal live-cell imaging of axon tips from DIV7 rat hippocampal neurons expressing mNG, mNG-KIF1A-WT, or mNG-KIF1A-T258M and stained with SiR-Lys. Single-channel images are shown in grayscale, and merged images in color with mNG or mNG-KIF1A in green and SiR- Lys in magenta. Arrowheads indicate representative mNG-KIF1A- and SiR-Lys-positive structures. Scale bars: 5 μm. **h,** Quantification showing that expression of mNG-KIF1A-T258M significantly increases SiR- Lys intensity at axon tips compared with mNG-KIF1A-WT. The graph shows individual data points and mean ± SD from 24–32 neurons per condition obtained from three independent cultures. Statistical significance was calculated by Kruskal–Wallis test followed by Dunn’s multiple-comparisons test. *P < 0.05; ****P < 0.0001; ns, not significant.

### KIF1A is required for lysosomal vesicle distribution in human neurons

Our observations in rat hippocampal neurons revealed an essential role for KIF1A in lysosomal vesicle transport. To determine whether KIF1A is similarly required for this process in human neurons, we employed iPSCs expressing the transcription factor Neurogenin-2 (NGN2) under the control of a doxycycline-inducible promoter.^64^ Upon doxycycline induction, these iPSCs undergo a well-defined differentiation program into glutamatergic neurons (i3Neurons) and form a mature AIS by 14 days of differentiation. This system therefore provides a useful platform for investigating the role of KIF1A in lysosomal vesicle transport during human neuronal development.

We first used CRISPR-Cas9 technology to delete the KIF1A gene in WT iPSCs (KIF1A-KO) and then reintroduced KIF1A into the KO line using a silencing-resistant piggyBac transposon-based system (Rescue) (Fig. S7a).^65,66^ WT, KIF1A-KO, and Rescue iPSCs were differentiated for 7 days, fixed, and immunostained for the somatodendritic marker MAP2 and the lysosomal marker LAMP1. Compared with WT i3Neurons, KIF1A-KO i3Neurons showed a marked reduction in LAMP1 signal in distal dendrites (Fig. 8a) and at axon tips (Fig. 8b). Similarly, KIF1A-KO i3Neurons showed a marked reduction in SiR-Lys staining in distal dendrites (Fig. 8c) and at axon tips (Fig. 8d). We performed similar analyses in 14-day-old i3Neurons, which have an AIS, and again observed reduced LAMP1 and SiR-Lys staining in dendrites (Fig. 8e,g) and at axon tips (Fig. 8f,h). These results are consistent with our observations in primary rat hippocampal neurons and further establish that KIF1A is required for lysosomal vesicle transport in both immature and mature neurons.

**Figure 8.**
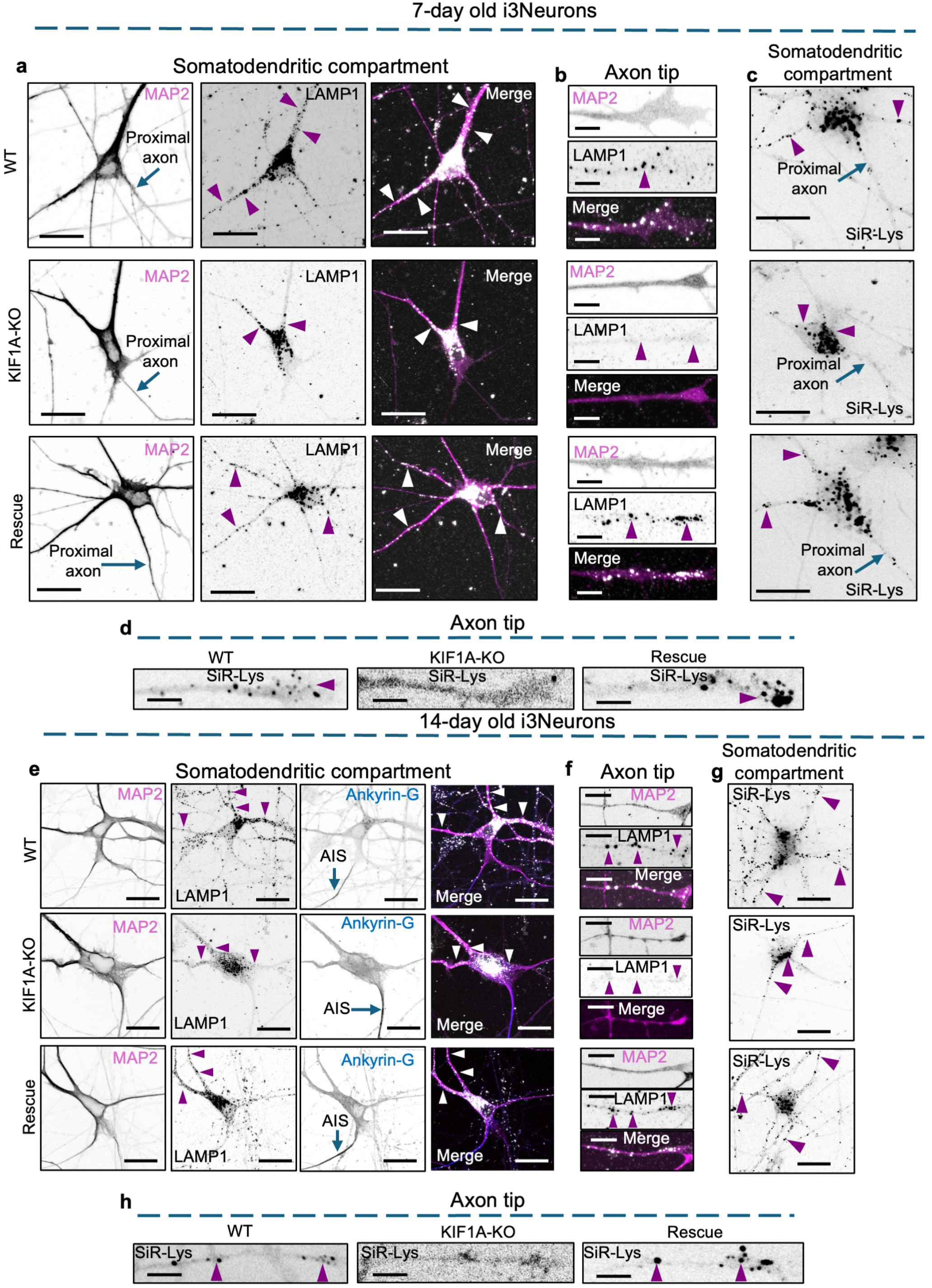
KIF1A is required for lysosomal vesicle distribution in human i3Neurons. **a,** Confocal fluorescence microscopy of the somatodendritic compartments of 7-day-old WT, KIF1A-KO, and KIF1A-KO i3Neurons rescued with human KIF1A (Rescue), fixed, permeabilized, and immunostained for LAMP1 and MAP2. The proximal axon is indicated. Single-channel images are shown in grayscale, and merged images in color with MAP2 in magenta and LAMP1 in white. Arrowheads indicate representative LAMP1-positive structures. Scale bars: 20 μm. **b,** Confocal fluorescence microscopy of axon tips from the 7-day-old i3Neuron lines described in a, immunostained for LAMP1 and MAP2. Arrowheads indicate representative LAMP1- positive structures. Scale bars: 5 μm. **c,** Confocal live-cell imaging of the somatodendritic compartments of 7-day-old WT, KIF1A-KO, and rescued i3Neurons stained with SiR-Lys. Arrowheads indicate representative SiR-Lys- positive structures. Scale bars: 20 μm. **d,** Confocal live-cell imaging of axon tips from 7-day-old WT, KIF1A-KO, and rescued i3Neurons stained with SiR-Lys. Arrowheads indicate representative SiR-Lys-positive structures. Scale bars: 5 μm. **e,** Confocal fluorescence microscopy of the somatodendritic compartments of 14-day-old WT, KIF1A-KO, and rescued i3Neurons fixed, permeabilized, and immunostained for LAMP1, MAP2, and ankyrin-G to identify the AIS. Single-channel images are shown in grayscale, and merged images in color with MAP2 in magenta, LAMP1 in white and Ankyrin-G in blue. Arrowheads indicate representative LAMP1-positive structures. Scale bars: 20 μm. **f,** Confocal fluorescence microscopy of axon tips from the 14-day-old i3Neuron lines described in e, immunostained for LAMP1 and MAP2. Arrowheads indicate representative LAMP1- positive structures. Scale bars: 5 μm. **g,** Confocal live-cell imaging of the somatodendritic compartments of 14-day-old WT, KIF1A- KO, and rescued i3Neurons stained with SiR-Lys. Arrowheads indicate representative SiR-Lys- positive structures. Scale bars: 20 μm. **h,** Confocal live-cell imaging of axon tips from 14-day-old WT, KIF1A-KO, and rescued i3Neurons stained with SiR-Lys. Arrowheads indicate representative SiR-Lys-positive structures. Scale bars: 5 μm.

Disruption of axonal lysosome transport has previously been shown to cause axonal swellings with aberrant accumulation of Tau and mitochondria.^67^ We observed a similar phenotype in i3Neurons differentiated from KIF1A-KO iPSCs, wherein they developed aberrant accumulation of Tau and mitochondria (Fig. S7b). Axonal lysosome transport has also been shown to be essential for mitochondrial health and function^67^ and loss of mitochondrial ΔΨm is known to elevate mitophagy.^68^ We immunostained KIF1A-KO i3Neurons and observed the presence of prominent LC3B puncta that co-localized with mitochondria (Fig. S7b), consistent with accumulation of mitophagy-related structures. Finally, i3Neurons differentiated from KIF1A-KO iPSCs also developed extensive axonal swellings (Fig. S7c). These observations thus establish the essential role of KIF1A in lysosomal vesicle transport in human neurons and demonstrate that loss of KIF1A is associated with profound defects in axonal integrity.

## Discussion

Lysosomal vesicles characterized by the presence of LAMP1 and other lysosomal proteins are distributed throughout neuronal dendrites and axons.^4,5,69^ This distribution requires long-range transport by molecular motors operating on a highly polarized and developmentally remodeled MT cytoskeleton. Previous work established a role for kinesin-1 (KIF5) motors in the axonal transport of lysosomal vesicles in non-neuronal cells and neurons.^20,21,70^ Here, we identify the kinesin-3 motor KIF1A as an essential motor for lysosomal vesicle transport in neurons and show that its ability to transport different lysosomal populations into the axon changes during neuronal maturation. We further identify the axon initial segment (AIS) as a site of this developmental regulation.

In immature rat hippocampal neurons, depletion of endogenous KIF1A reduces the abundance of LAMP1-positive and SiR-Lys-positive lysosomal vesicles in dendrites and axons, whereas KIF1A overexpression promotes the accumulation of both populations at distal dendrites and axon tips. KIF1A is therefore required for lysosomal vesicle distribution during early neuronal development. KIF1A depletion from mature rat hippocampal neurons also reduced both LAMP1 and SiR-Lys in dendrites and at axon tips. KIF1A overexpression, however, increased axonal LAMP1 accumulation without increasing SiR-Lys accumulation. This difference reflects a developmental change in the lysosomal cargo transported by KIF1A into the axon. In DIV3 neurons, a substantial fraction of anterogradely moving LAMP1-positive vesicles corresponded to SiR-Lys-positive, and KIF1A overexpression increased the number of SiR-Lys-positive tracks entering the axon. At the mature AIS, a smaller proportion of anterograde LAMP1-positive or KIF1A-positive tracks colocalized with SiR-Lys, SiR-Lys tracks were less bright, and increasing KIF1A expression no longer increased SiR-Lys-positive axonal entry. Thus, KIF1A remains capable of transporting lysosomal vesicles into mature axons but preferentially transports a less degradative population after AIS formation.

The requirement for KIF1A also raises the question of how its function relates to that of other kinesin motors expressed in neurons. KIF1A is a neuronally enriched member of the kinesin-3 family,^24^ whereas the closely related KIF1B is more widely expressed and is also present in neurons.^71^ Neurons additionally express the kinesin-1 motors KIF5A, KIF5B, and KIF5C, which have established roles in axonal transport, including lysosomal vesicle transport.^20,21,70^ Nevertheless, these motors cannot fully compensate for KIF1A loss, since KIF1A depletion disrupts lysosomal vesicle distribution despite their presence. This specificity could reflect preferential association of different kinesins with lysosomal vesicles that differ in composition, maturation state, or destination. Alternatively, different motors could act at distinct steps of a common transport pathway, making more than one kinesin necessary for efficient delivery from the soma to distal axonal regions. Distinguishing among these possibilities will require determining which motors associate with defined lysosomal populations and where along the transport pathway they act.

The ability of overexpressed KIF1A to increase LAMP1 and synaptophysin accumulation at mature axon tips argues against a general inhibition of KIF1A-mediated axonal transport after AIS formation. Instead, the AIS appears to regulate KIF1A in a cargo-dependent manner. AIS MTs adopt a lattice conformation resembling the GTP-bound state^54^ and are enriched in the plus-end-binding proteins EB1 and EB3 through their interaction with AnkG.^55,56^ These features are characteristic of dynamic MTs and distinguish the AIS MT environment from more stable regions of the neuronal MT cytoskeleton. KIF1A has reduced affinity for GTP-tubulin-rich MTs, and EB3 further reduces KIF1A–MT binding.^57^ The MT environment that develops at the mature AIS could therefore regulate KIF1A engagement with the lattice. Importantly, this environment does not generally prevent KIF1A-mediated cargo entry into the axon. Instead, it preferentially restricts entry of more degradative lysosomal vesicles while permitting transport of less degradative lysosomal vesicles and other KIF1A cargoes.

The effects of AnkG and EB depletion directly support this interpretation. Disrupting EB- protein enrichment, either indirectly by AnkG depletion or directly by EB1+EB3 depletion, allowed KIF1A to transport more and brighter SiR-Lys-positive lysosomal vesicles into mature axons without increasing overall anterograde KIF1A traffic. How altered KIF1A–MT interactions produce this cargo selectivity remains unclear. KIF1A is highly processive under low-load conditions,^72^ whereas under hindering load it undergoes rapid detachment but is kinetically tuned for repeated reattachment to the MT lattice.^73^ Anterogradely transported axonal LAMP1-positive carriers include relatively small, weakly acidic vesicles.^7^ Differences in the physical properties of lysosomal vesicle populations could therefore impose different loads on KIF1A. Reduced KIF1A–MT affinity at the EB-rich AIS might be sufficient for movement of relatively low-load lysosomal carriers but become limiting for transport of larger, more mature, catalytically active lysosomes. Although this possibility remains to be tested directly, it could explain why disrupting EB proteins changes the lysosomal cargo transported through the AIS without increasing KIF1A traffic itself.

The KIF1A-T258M mutant provides a complementary test from the motor side. T258M is associated with hereditary spastic paraplegia type 30 and exhibits increased affinity for GTP- tubulin.^57,63,74^ Consistent with the proposed role of the AIS MT environment, KIF1A-T258M transported more SiR-Lys-positive cargo through the GTP-tubulin-like AIS lattice than KIF1A- WT, despite not increasing the overall number of anterograde KIF1A tracks. Thus, perturbing either the AIS MT environment or the ability of KIF1A to interact with that environment produces a similar change in lysosomal cargo transport.

What are the SiR-Lys-positive vesicles that gain increased access to the axon under these conditions? Mature, degradative lysosomes are enriched in the neuronal soma, and some soma-derived lysosomes containing active cathepsins undergo anterograde transport into distal axons, where they contribute to local degradative capacity.^75^ The increased number and intensity of anterogradely moving SiR-Lys-positive vesicles following AnkG or EB1+EB3 depletion or expression of KIF1A-T258M may therefore represent increased KIF1A-dependent entry of mature, more degradative lysosomes from the soma. The mature AIS would thus not completely exclude degradative lysosomes from the axon but would restrict their KIF1A- dependent entry relative to that of less degradative lysosomal vesicles.

A related form of spatial regulation of KIF1A occurs at presynaptic sites, where dynamic, GTP-tubulin-rich MTs and EB proteins weaken KIF1A–MT interactions and promote local unloading of SVPs.^57^ At the AIS, similar properties of the MT lattice appear to serve a different function by restricting passage of specific KIF1A-associated lysosomal cargo into the mature axon. Reduced KIF1A affinity for GTP-tubulin- and EB-rich MTs may therefore provide a common mechanism by which distinct MT domains locally control KIF1A cargo transport. Consistent with this idea, KIF1-family motors are also locally regulated at axonal branch junctions, where cargo transport is selectively directed toward specific branches.^76^ Dynein regulators at the AIS similarly contribute to selective retrograde cargo transport.^77^ Local regulation of both anterograde and retrograde motors may therefore contribute to selective cargo passage and distribution within axons.

The behavior of KIF1A-T258M also suggests a mechanism by which disease-associated KIF1A mutations could alter neuronal transport beyond simply increasing or decreasing motor activity.^74,78,79^ Altered motor–MT affinity could additionally change where particular cargoes are transported by allowing them to cross transport boundaries that normally restrict their distribution. Whether altered lysosomal vesicle distribution contributes to neuronal dysfunction associated with T258M or other KIF1A mutations remains to be determined.

The requirement for KIF1A in lysosomal vesicle distribution extends to human iPSC-derived i3Neurons. KIF1A disruption reduced LAMP1- and SiR-Lys-positive structures in distal dendrites and at axon tips in both immature and mature i3Neurons, and these phenotypes were restored by KIF1A re-expression. Because i3Neurons could not be maintained at sufficiently low density for reliable visualization of individual AISs, we could not directly test developmental regulation of KIF1A-driven lysosome transport through the AIS. KIF1A loss was also associated with extensive axonal swellings containing abnormal accumulations of Tau and mitochondria, together with LC3B- and mitochondria-positive structures. Similar axonal swellings, mitochondrial defects, and accumulation of autophagic structures occur following disruption of axonal lysosomal vesicle transport.^67^ The phenotypes observed in KIF1A-deficient i3Neurons are therefore consistent with an essential role for KIF1A-dependent lysosomal vesicle distribution in maintaining human axonal integrity.

In summary, KIF1A is an essential motor for neuronal lysosomal vesicle transport whose cargo-transporting activity changes during neuronal maturation. Following AIS formation, the local MT environment preferentially restricts KIF1A-dependent entry of lysosomal vesicles containing detectable cathepsin D activity while permitting transport of less degradative lysosomal cargo. Manipulating either the AIS MT environment or KIF1A interaction with this environment relaxes this restriction and increases the entry of more degradative lysosomal vesicles into mature axons. The AIS therefore functions as a developmental regulator of KIF1A- driven lysosomal transport, illustrating how local differences in motor–MT interactions can determine cargo distribution in neurons.

## Materials and methods

All animal experiments were performed in accordance with National Institutes of Health and U.S. government regulations. Research was conducted under Animal Study Proposals 24-021 and 25-011 approved by the *Eunice Kennedy Shriver* National Institute of Child Health and Human Development Animal Care and Use Committee.

### Antibodies and reagents

The following antibodies were used for immunofluorescence microscopy at the indicated dilutions: chicken-anti-MAP2 (Abcam, Cat. ab5392, 1:2,000); mouse anti-rat LAMP1 (6H2, generated in-house, 1:2 culture supernatant); goat anti-ankyrin G (Santa Cruz Biotechnology, Cat. sc-31778, 1:300); chicken anti-GFP (Invitrogen, Cat. A10262, 1:2,000); rabbit anti-SYP (Abcam, Cat. ab32127, 1:2,000); rat anti-mCherry (Invitrogen, Cat. M11217, 1:2,000); mouse anti- EB1 (BD Biosciences, Cat. 610535, 1:300); mouse anti-LAMP1 (DSHB, Cat. H4A3, 1:500); rabbit anti-TOM20 (Proteintech, Cat. 11802-1-AP, 1:1,000); mouse anti-Tau-1 (clone PC1C6; human reactivity) (Millipore Sigma, Cat. MAB3420,1:500); mouse anti-CYCS (6H2.B4) (BD Biosciences Cat#556432, 1:200); rabbit anti-LC3 (D11) (Cell Signaling Technology, Cat#3868, 1:200); donkey anti-rabbit Alexa Fluor 488 (Invitrogen, Cat. A-21206, 1:1,000); donkey anti-rabbit Alexa Fluor 555 (Invitrogen, Cat. A-31572, 1:1,000); donkey anti-rabbit Alexa Fluor 647 (Invitrogen, Cat. A- 31573, 1:1,000); donkey anti-mouse Alexa Fluor 488 (Invitrogen, Cat. A-21202, 1:1,000); donkey anti-mouse Alexa Fluor 555 (Invitrogen, Cat. A-31570, 1:1,000); donkey anti-mouse Alexa Fluor 647 (Invitrogen, Cat. A-31571, 1:1,000); donkey anti-goat Alexa Fluor 488 (Invitrogen, Cat. A- 11055, 1:1,000); donkey anti-goat Alexa Fluor 647 (Invitrogen, Cat. A-21447, 1:1,000); donkey anti-chicken Alexa Fluor 488 (Invitrogen, Cat. A78948, 1:1,000); donkey anti-chicken Alexa Fluor 647 (Invitrogen, Cat. A78952, 1:1,000); donkey anti-rat Alexa Fluor 555 (Invitrogen, Cat. A48270, 1:1,000).

The following antibodies were used for immunoblotting at the indicated dilutions: rabbit anti- KIF1A (Abcam, Cat. ab180153, 1:1,000), goat anti-rabbit-HRP (Perkin Elmer, Cat. NEF812001EA, 1:3,000), mouse anti-α-tubulin–HRP (Santa Cruz Biotechnology, Cat. sc-32293, 1:3,000), goat anti-mouse-HRP (Perkin Elmer, Cat. NEF822001EA, 1:3,000).

### Expression constructs and shRNA

peGFP-N1 was obtained from Clontech. The eGFP-mKIF1A was previously described in Guardia et al., 2016 (Addgene Cat. 172206). The mNeonGreen-mKIF1A construct was made by cloning the mKIF1A sequence from eGFP-mKIF1A into the mNeonGreen-C1 backbone (Addgene Cat. 197924^80^). The LAMP1-mNeonGreen construct was made by cloning the mNeonGreen sequence into the LAMP1-GFP^21^ plasmid. The mNeonGreen-mKIF1A-T258M plasmid was prepared by introducing a point mutation in the mNeonGreen-mKIF1A-WT plasmid using Q5 Site-Directed Mutagenesis Kit (E0554, New England Biolabs). To rescue KIF1A-KO cells, iPSCs were transfected with a silencing-resistant piggyBac plasmid (M54) that contained mNeonGreen-hKIF1A-WT sequence cloned between the 5’ and 3’ inverted terminal repeats (ITRs). The M54 piggyBac plasmid was a gift from Dr. Michael E. Ward (NINDS, NIH).

All shRNA plasmid constructs consisted of the target sequence (under the U6 promoter) followed by a BFP, GFP or mCherry sequence (under the hPGK promoter). In all experiments, the scrambled shRNA sequence used was 5’-gatgaacctgatgacgttc-3’. For KIF1A KD, the targeted sequence was 5’-tacctatgtgaacggcaagaa-3’.^61^ For AnkG KD, two AnkG sequences were targeted: (1) 5’- gctgtttgcaacttaaatatt-3’; (2) 5’-ccacaacagctcgccttattt-3’. For EB1 plus EB3 KD, the targeted sequences were EB1 5’-tctgacaaagatagaacag-3’ and EB3 5’- tatgatggaaaggattac-3’.^81^

KIF1A-shRNA KD efficacy was verified by immunoblotting cell lysates of rat hippocampal neurons infected with lentiviral particles on DIV3 and lysed on DIV7. Lentiviral particles expressing scrambled shRNA or KIF1A shRNA were generated in HEK293T cells by co-transfecting scrambled/KIF1A shRNA transfer plasmids with the packaging plasmids psPAX2 (Addgene, cat. 12260), pMD2.G (Addgene, Cat. 12259), and pAdVantage (Promega, Cat. E1711). Viral supernatants from the HEK293T cell culture were collected, filtered, concentrated and added to hippocampal neuron culture at DIV3 and the cells were lysed on DIV7 for SDS-PAGE and immunoblotting.

### Rat hippocampal neuron culture

Primary hippocampal cultures were prepared from embryonic day 18 (E18) rat brains as described previously.^82,83^ Dissociated hippocampal neuronal cells were plated on 18mm coverslips, inserted into wells of a 12-well plate and pre-treated with poly-L-lysine hydrobromide ( Sigma-Aldrich, p2636) and laminin (Roche, 11243217001) at a density of 40,000 cells/coverslip. Hippocampal neurons were cultured in complete Neurobasal medium (CNB) consisting of Neurobasal medium (NB) (Gibco, 21103049), supplemented with 1X B27, serum free (Gibco, 17504044), 2mM L-glutamine (Quality Biological, 118-084-671), and 100 U ml^−1^ penicillin–streptomycin (Quality Biological, 50-751-7267).

### Transfection

Rat hippocampal neurons were transfected using Lipofectamine 2000 (Invitrogen, 11668027). For each 18-mm coverslip in a well of a 12-well plate, the following procedure was used: 0.7-1 μg of total plasmid DNA was added to 75 μL incomplete NB medium and the tube was flicked gently to mix the contents. Separately, 1.5 μL of Lipofectamine 2000 was mixed in 75 μL incomplete NB medium. The DNA and Lipofectamine mixtures were combined, mixed gently and incubated for 20 min. While the transfection mix was incubating, 700 μL of medium was aspirated from each well to be transfected, stored as conditioned medium, and replaced with 700 μL of prewarmed (37°C) incomplete NB medium. The transfection mix was added to the wells and gently mixed, and the cells were transferred to a 37°C incubator for 1 h. Then the medium in each transfected well was replaced with prewarmed (37°C) incomplete NB medium; this step was repeated twice. Finally, the conditioned medium was added back to the wells and topped up with freshly prepared CNB to 1 mL/well. Throughout transfection, the cells were kept in at least ∼200 μL of medium and were not exposed to air.

### Immunofluorescence microscopy

For immunofluorescence experiments, neurons grown on coverslips were first fixed with 4% paraformaldehyde (PFA) (Thermo Scientific, 28906) in 1× PBS at room temperature for 20 min. Neurons were then washed twice in 1× PBS and then incubated for 1 h at room temperature in immunofluorescence (IF) buffer (1× PBS, 1% BSA and 0.1% saponin). Neurons were then incubated in IF buffer with appropriate primary antibodies either for 1 h at 37°C or overnight at 4°C. After primary antibody incubation, neurons were washed in fresh IF buffer three times for 5 min each. Then, the coverslips were incubated in IF buffer with secondary antibodies for 45 min. Coverslips were washed in 1× PBS twice for 10 min each and mounted on slides. Confocal images were acquired using a Zeiss LSM 880 or LSM 900 microscope with a 63× oil-immersion objective.

### Live-cell imaging

Live-cell imaging of rat hippocampal neurons and i3Neurons was performed on a spinning-disk microscope system comprising a Nikon Eclipse Ti-E inverted research microscope equipped with a Perfect Focusing System, Plan Apo VC 60× 1.4 NA oil objective (Nikon), CSU- X1 A1 spinning disk (Yokogawa), Hamamatsu 4ORCA-Flash4 CMOS camera, and Nikon Laser Stack controlled by a camera trigger. All videos were acquired with 75-ms exposure per image and appropriate laser intensity. Single-channel videos were acquired with a 100-ms interval (10 frames per second or fps), and dual-channel videos were acquired with no delay (∼6.66 fps). During live-cell imaging, cells were maintained at 37°C and 5% CO₂ in a stage-top incubator (Tokai HIT INU). For live-cell imaging, lysosomal vesicles were labeled using SiR-Lysosome (Cytoskeleton Inc., Cat. CY-SC012, 1:1,000). The AIS of DIV7 live rat hippocampal neurons was labeled using mouse anti-pan-neurofascin (NF) extracellular antibody (Antibodies Inc., Cat. 75- 172) conjugated with CF555 (Biotium, Cat. 92234) and used at a 1:1,000 dilution in NB medium.

### Human iPSC and i3Neuron culture

Human iPSCs expressing doxycycline-inducible *NGN2* (neurogenin)^64,84^ were cultured on Matrigel-coated dishes in Essential 8 Flex Medium (E8Flex) (Gibco, Cat. A2858501) supplemented with 10 μM Y-27632 dihydrochloride ROCK inhibitor (Selleck Chemicals, Cat.S1049). Differentiation of iPSCs to i3Neurons was performed as described previously.^64,67,84^ Briefly, iPSCs were seeded on day 0 in induction medium containing DMEM/F12 (Gibco, Cat. 11330032) with 1× non-essential amino acids (NEAAs) (Gibco, Cat. 11140050), 1× GlutaMAX (Gibco, Cat. 35050061), 1× N-2 supplement (Gibco, Cat. 17502048), 2 μg ml^−1^ doxycycline (Cat. D9891, Sigma-Aldrich), and 10 μM Y-27632 dihydrochloride (Rho kinase inhibitor). After 48 h, the induction medium was replaced with complete BrainPhys (CBP) neuronal culture medium consisting of BrainPhys medium (STEMCELL Technologies, Cat. 5790) supplemented with 10 ng ml^−1^ neurotrophin-3 (PeproTech, Cat. 450-03), 10 ng ml^−1^ BDNF (PeproTech, Cat. 450-02), 1× serum-free B-27 supplement (Gibco, Cat. 17504044), 2 μg ml^−1^ doxycycline and 1 μg ml^−1^ mouse laminin (Gibco, Cat. 23017015). Twenty-four h later, the cells were detached using Accutase (STEMCELL Technologies, Cat, 07920), resuspended in CBP and plated on 18- mm coverslips (one per well of a 12-well plate) pre-treated with poly-L-lysine hydrobromide and laminin. Neurons were cultured in CBP medium and processed for immunofluorescence microscopy or live-cell imaging on day 7 or 14.

### CRISPR-Cas9 KO of KIF1A

To disrupt the *KIF1A* gene in iPSCs, we used the CRISPR-Cas9 system. Using CRISPRdirect (https://crispr.dbcls.jp/), we identified two sequences within exon 2 of the human *KIF1A* gene as sites for Cas9-mediated scission. Next, the gRNA target sequences [indicated in UPPER CASE] complementary to these sequences were incorporated into oligonucleotides with overhangs [indicated in lower case] to enable cloning into the PX458 plasmid (gift from F. Zhang; Addgene Cat. 48138). The oligonucleotide pairs used for cloning were: (1) 5’- caccGGCCGGGGCTTCGGTGAAGG-3’ and 5’-aaacCCTTCACCGAAGCCCCGGCC-3’, and (2) 5’-caccTGGAGTCACGGCTCATTTCC-3 and 5’-aaacGGAAATGAGCCGTGACTCCA-3’. Each of the oligonucleotide pairs was annealed and cloned separately into the PX458 plasmid. Insertion was verified by sequencing. The two gRNA-containing plasmids were transfected into iPSCs using Lipofectamine 3000 (Invitrogen, Cat. L3000001). After 48 h, single iPSCs were sorted into a 96-well plate based on positive GFP signal. The cells were cultured in E8 Flex Medium with RevitaCell Supplement (Gibco, Cat. A2644501) for 14 days. The plate was visually scanned for colonies under a brightfield microscope and healthy-looking, large colonies were lifted using Accutase. The resuspended cells from each colony were cultured separately and expanded until confluent in a 60-mm TC plate. Immunoblotting was performed to verify KIF1A gene inactivation in each of the cultures. One of the clones was selected for further experiments.

### Rescue of KO cells

To rescue KIF1A-KO cells, we used a silencing-resistant piggyBac plasmid backbone (M54, Gift from Dr. Michael Ward, NINDS, NIH)^66^ containing the mNeonGreen–human KIF1A transposon insert. Briefly, the KIF1A-KO iPSCs were co-transfected with M54-mNeon-hKIF1A and a transposase plasmid pBase-mCherry (Vector Builder) using Lipofectamine-3000. After 48 h, the growth medium was supplemented with 0.5 μg/mL puromycin (GoldBio, Cat. P-600-100). After 24 h, the remaining cells were cultured for a further 10 days in E8 Flex Medium containing 0.5 μg/mL puromycin with regular medium changes and replating as needed. Stable expression of mNeonGreen-KIF1A was verified by immunoblotting.

### Image analysis, quantification and statistical analysis

Confocal images of immunofluorescence samples were processed and quantified using Fiji. Prior to quantification, confocal z-stacks were converted to maximum-intensity projections. To quantify lysosome accumulation at dendritic and axon tips, a 20-pixel-wide, 15–30-μm-long line was drawn over the region of interest, and the mean fluorescence intensity was measured. For intensity measurements in Figures 1–4, values obtained from dendrites represent the mean fluorescence intensity measured from at least three or four dendrites per cell. For quantification of lysosomal puncta area within the somatodendritic compartment, fluorescence images were thresholded in Fiji, the area of individual puncta was measured, and the mean punctum area was calculated for each cell.

Live-cell videos acquired using spinning-disk fluorescence microscopy were processed using Fiji. KIF1A–SiR-Lys colocalization within the proximal axon or AIS was quantified from kymographs generated separately for each channel using the Reslice function in Fiji without interpolation. SiR-Lys-positive anterograde traces were first marked and added to the ROI Manager. These ROIs were then applied to the corresponding KIF1A kymograph to identify SiR-Lys traces that colocalized with KIF1A. Additional KIF1A-positive traces lacking detectable SiR-Lys signal were then added to the ROI Manager to determine the total number of KIF1A traces. The proportion of KIF1A traces colocalizing with SiR-Lys was calculated as the number of KIF1A/SiR-Lys-colocalizing traces divided by the total number of KIF1A traces. The same approach was used to quantify the colocalization between LAMP1 and SiR-Lys movement through the proximal axon/AIS. Anterograde track numbers in kymographs were determined by visual counting after adjusting image contrast to ensure that faint tracks were also detected. To quantify the fluorescence intensity of moving SiR-Lys-positive tracks, a 5-pixel-wide line was drawn along each track in the kymograph, avoiding overlap with neighboring tracks as much as possible, and mean fluorescence intensity was measured. KIF1A track velocity was calculated from the slope of individual tracks in the kymograph by drawing a line along the track and converting the measured slope to velocity using the spatial and temporal calibration of the image.

Measured intensities were tabulated in Microsoft Excel and statistical analyses were performed using GraphPad Prism 11. Bar graphs show the mean ± SD. The total number of cells analyzed is specified in the corresponding figure legends. Data sets were tested for normality by Kolmogorov-Smirnov (KS) test, and statistical differences were assessed using appropriate parametric or non-parametric tests as indicated in the corresponding figure legends. In all the data sets the analysis unit was the individual neuron and the *P* < 0.05 was considered significant.

### Immunoblotting

Neurons were washed with 1× PBS and lysed in 1× Laemmli buffer (Bio-Rad, Cat. 1610747) supplemented with 5% 2-mercaptoethanol (Millipore Sigma, Cat. M3148) for 5 min at room temperature. Lysates were boiled at 95°C for 5 min and subjected to SDS-PAGE. Proteins from polyacrylamide gels were then transferred onto nitrocellulose membranes, which were then blocked for 30 min in blocking buffer containing 5% nonfat milk prepared in Tris-buffered saline with 0.01% Tween-20 (TBST; Millipore Sigma, Cat. P1379). Membranes were incubated overnight at 4 °C with primary antibodies diluted in blocking buffer with gentle rocking. After primary antibody incubation blots were washed twice (10 min each) in 1x TBST and incubated for 30 min with the appropriate secondary antibodies diluted in blocking buffer. After three additional washes in TBST, the immunoblots were developed using Clarity Western ECL substrate (Bio-Rad, Cat. 1705061) and imaged using a ChemiDoc MP imaging system (Bio-Rad, Cat. 12003154).

## Data availability

Data are available in the article and its supplementary materials, or otherwise available upon request.

## Acknowledgements

We thank Xiaolin Zhu for help in the preparation of rat embryonic primary hippocampal neurons, and Michael E. Ward for the kind gift of reagents. This work was supported by the Intramural Program of the NICHD (project ZIA HD001607) and a NICHD FY25 Career Development Award to N.A. Tirumala.

## Author contributions

Conceptualization: N. A. Tirumala and J. S. Bonifacino; Methodology: N. A. Tirumala; Investigation: N. A. Tirumala, Neuron culture: N.A Tirumala and Surbhi Verma; Writing: N. A. Tirumala and J. S. Bonifacino; Funding acquisition: J. S. Bonifacino and N.A. Tirumala; Supervision: J. S. Bonifacino.

## Supplementary material

**Figure S1.**
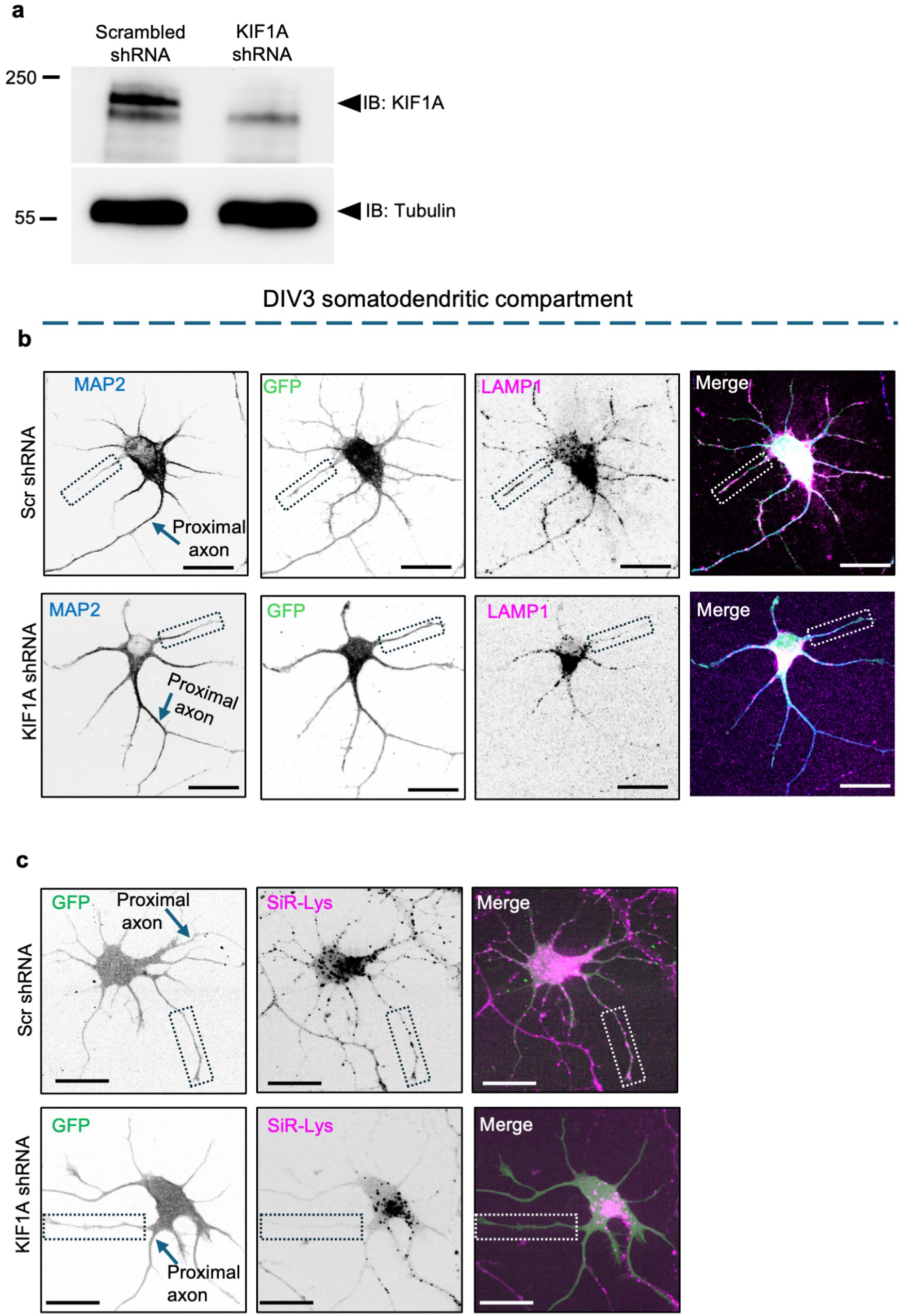
KIF1A depletion reduces lysosomal vesicle localization in the dendrites and axons of immature rat hippocampal neurons. **a,** Immunoblot showing depletion of KIF1A in rat hippocampal neurons infected with lentivirus encoding KIF1A shRNA. Tubulin was used as a loading control. Molecular masses are indicated in kilodaltons. **b,** Confocal fluorescence microscopy of the somatodendritic compartment of DIV3 rat hippocampal neurons expressing Scr shRNA or KIF1A shRNA along with GFP, fixed, permeabilized, and immunostained for MAP2 and LAMP1. The dashed outlines indicate the dendrite tips shown in Fig. 1b. Single-channel images are shown in grayscale, and merged images in color with MAP2 in blue, GFP in green, and LAMP1 in magenta. Scale bars: 20 μm. **c,** Confocal live-cell imaging of the somatodendritic compartment of DIV3 rat hippocampal neurons expressing Scr shRNA or KIF1A shRNA along with GFP and stained with SiR-Lys. The dashed outlines indicate the dendrite tips shown in Fig. 1f. Single-channel images are shown in grayscale, and merged images in color with GFP in green and SiR-Lys in magenta. Scale bars: 20 μm.

**Figure S2.**
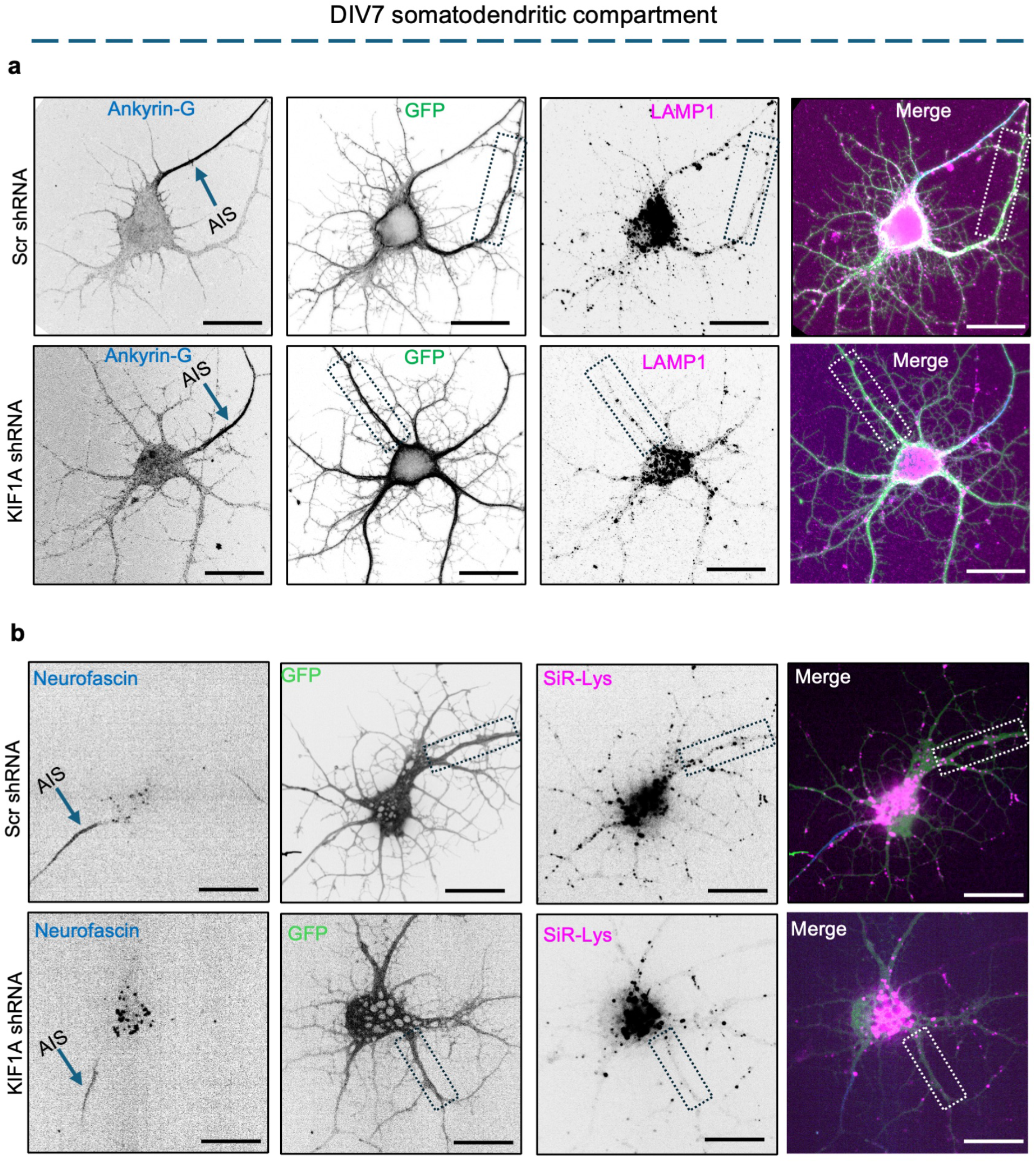
KIF1A depletion reduces lysosomal vesicle localization in the dendrites and axons of mature rat hippocampal neurons. **a,** Confocal fluorescence microscopy of the somatodendritic compartment of DIV7 rat hippocampal neurons expressing Scr shRNA or KIF1A shRNA along with GFP, fixed, permeabilize, and immunostained for ankyrin-G and LAMP1. The dashed outlines indicate the dendrites shown in Fig. 2a. Single-channel images are shown in grayscale, and merged images in color with ankyrin-G in blue, GFP in green, and LAMP1 in magenta. Scale bars: 20 μm. **b,** Confocal live-cell imaging of the somatodendritic compartment of DIV7 rat hippocampal neurons expressing Scr shRNA or KIF1A shRNA along with GFP, stained with SiR-Lys and labeled for neurofascin to identify the AIS. The dashed outlines indicate the dendrites shown in Fig. 2e. Single-channel images are shown in grayscale, and merged images in color with neurofascin in blue, GFP in green, and SiR-Lys in magenta. Scale bars: 20 μm.

**Figure S3.**
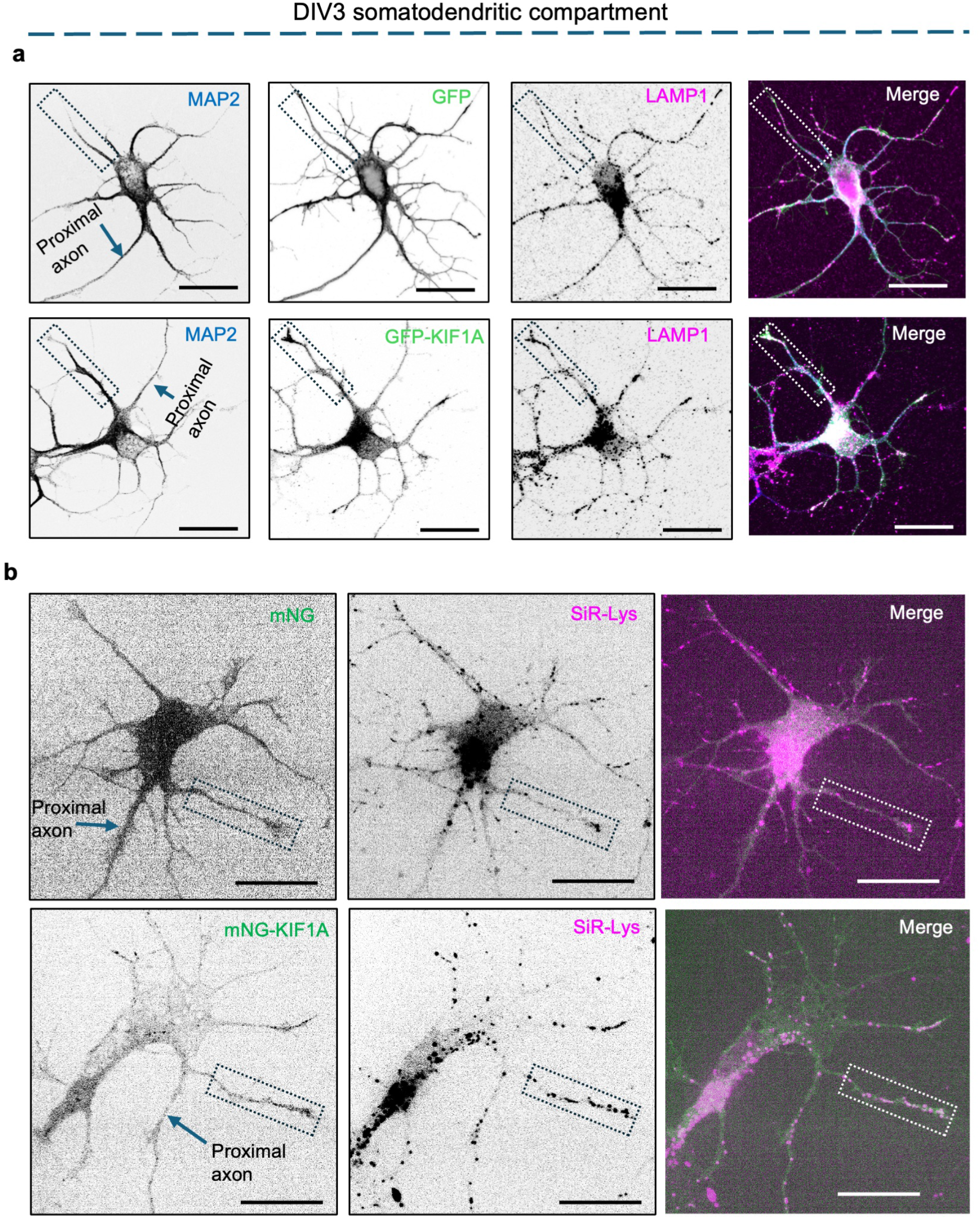
KIF1A expression promotes lysosomal vesicle accumulation in the dendrites and axons of immature rat hippocampal neurons. **a,** Confocal fluorescence microscopy of the somatodendritic compartment of DIV3 rat hippocampal neurons expressing GFP or GFP-KIF1A, fixed, permeabilized, and immunostained for MAP2 and LAMP1. The dashed outlines indicate the dendrite tips shown in Fig. 3a. Single-channel images are shown in grayscale, and merged images in color with MAP2 in blue, GFP or GFP-KIF1A in green, and LAMP1 in magenta. Scale bars: 20 μm. **b,** Confocal live-cell imaging of the somatodendritic compartment of DIV3 rat hippocampal neurons expressing mNG or mNG-KIF1A and stained with SiR-Lys. The dashed outlines indicate the dendrite tips shown in Fig. 3e. Single-channel images are shown in grayscale, and merged images in color with mNG or mNG-KIF1A in green and SiR-Lys in magenta. Scale bars: 20 μm.

**Figure S4.**
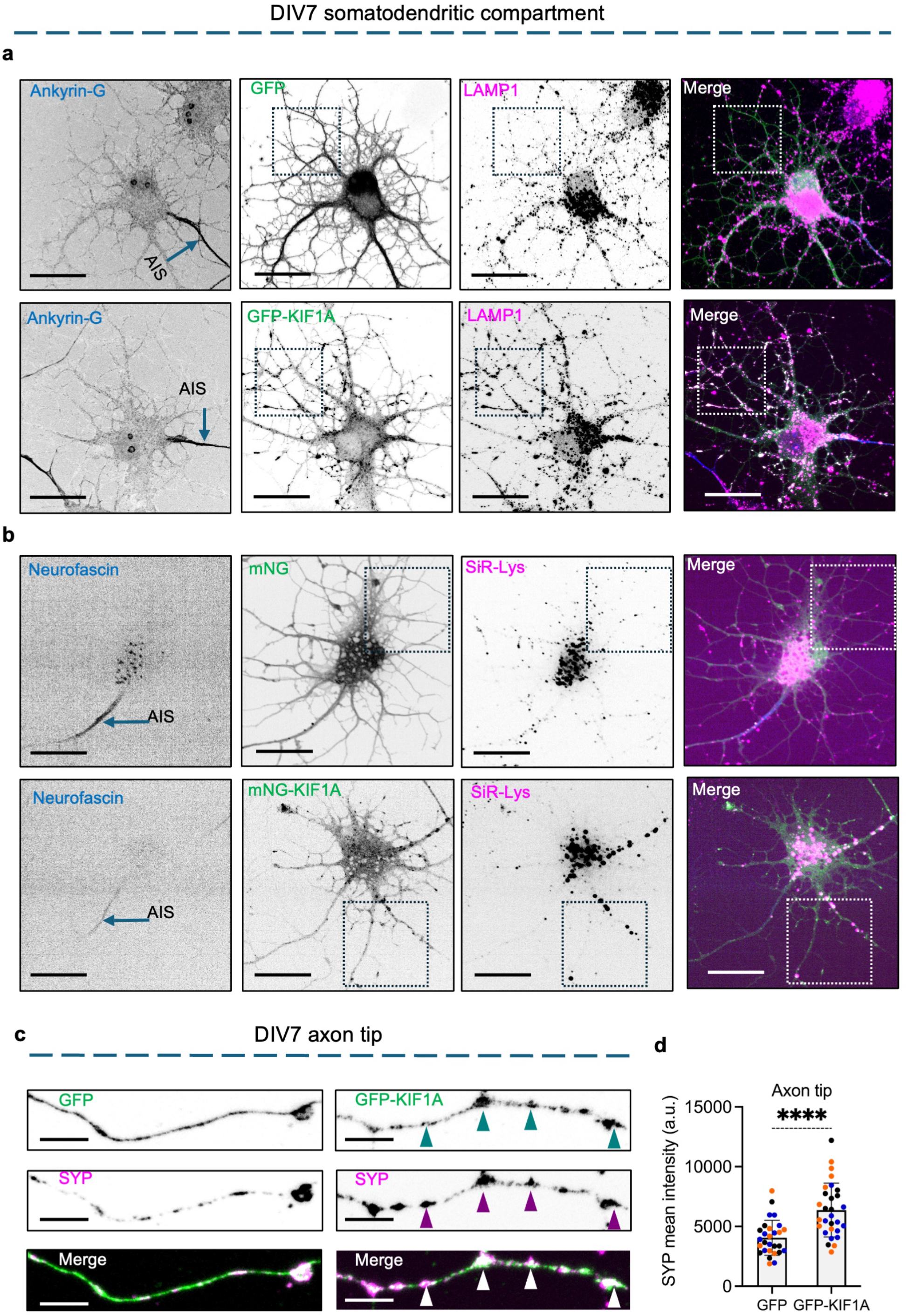
In mature rat hippocampal neurons, KIF1A expression increases accumulation of less degradative lysosomes and synaptophysin at the axon tips. **a,** Confocal fluorescence microscopy of the somatodendritic compartment of DIV7 rat hippocampal neurons expressing GFP or GFP-KIF1A, fixed, permeabilized, and immunostained for ankyrin-G and LAMP1. The dashed outlines indicate the dendritic regions shown in Fig. 4a. Single-channel images are shown in grayscale, and merged images in color with ankyrin-G in blue, GFP or GFP-KIF1A in green, and LAMP1 in magenta. Scale bars: 20 μm. **b,** Confocal live-cell imaging of the somatodendritic compartment of DIV7 rat hippocampal neurons expressing mNG or mNG-KIF1A, stained with SiR-Lys and labeled for neurofascin to identify the AIS. The dashed outlines indicate the dendritic regions shown in Fig. 4e. Single-channel images are shown in grayscale, and merged images in color with neurofascin in blue, mNG or mNG-KIF1A in green, and SiR-Lys in magenta. Scale bars: 20 μm. **c,** Confocal fluorescence microscopy of axon tips from DIV7 rat hippocampal neurons expressing GFP or GFP-KIF1A, fixed, permeabilized, and immunostained for synaptophysin (SYP). Single-channel images are shown in grayscale, and merged images in color with GFP or GFP-KIF1A in green and SYP in magenta. Arrowheads indicate representative GFP-KIF1A- and SYP-positive structures. Scale bars: 5 μm. **d,** Quantification showing that GFP-KIF1A expression significantly increases SYP intensity at the axon tip. The graph shows individual data points and mean ± SD from 28–29 neurons per condition obtained from three independent cultures. Statistical significance was calculated by two-tailed unpaired Welch’s t-test. ****P < 0.0001.

**Figure S5.**
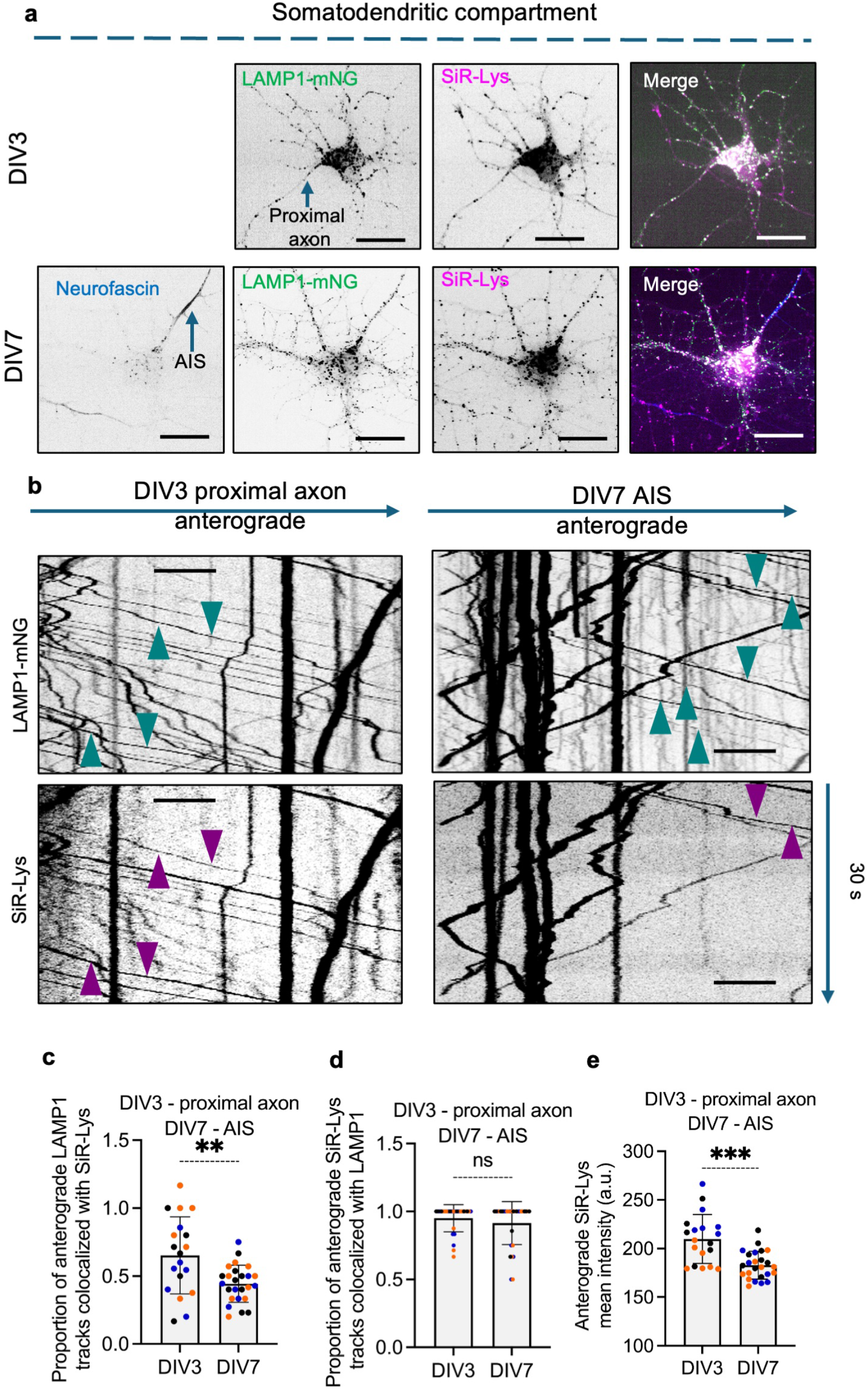
Neuronal maturation reduces the anterograde transport of degradative LAMP1-positive lysosomal vesicles. **a,** Confocal live-cell imaging of the somatodendritic compartments of DIV3 and DIV7 rat hippocampal neurons expressing LAMP1-mNeonGreen (LAMP1-mNG) and stained with SiR-Lys. Neurofascin labeling identifies the AIS in DIV7 neurons. Single-channel images are shown in grayscale, and merged images in color with LAMP1-mNG in green, SiR-Lys in magenta, and neurofascin in blue where shown. Scale bars: 20 μm. **b,** Kymographs generated from live videos of the proximal axon of DIV3 neurons and the AIS of DIV7 neurons expressing LAMP1-mNG and stained with SiR-Lys. Teal arrowheads indicate representative LAMP1-mNG-positive anterograde tracks, and magenta arrowheads indicate SiR-Lys- positive anterograde tracks. Scale bars: 5 μm. **c,** Quantification showing that the proportion of anterograde LAMP1-mNG-positive tracks colocalizing with SiR-Lys is significantly lower at the AIS of DIV7 neurons than in the proximal axon of DIV3 neurons. The graph shows individual data points and mean ± SD from 19–25 neurons per condition obtained from three independent cultures. Statistical significance was calculated by two-tailed Mann–Whitney test. **P < 0.01 **d,** Quantification showing the proportion of anterograde SiR-Lys-positive tracks colocalizing with LAMP1-mNG in DIV3 and DIV7 neurons. The proportion is close to one in both developmental stages and is not significantly different between DIV3 and DIV7 neurons. The graph shows individual data points and mean ± SD from 19–25 neurons per condition obtained from three independent cultures. Statistical significance was calculated by two-tailed Mann–Whitney test. ns, not significant. **e,** Quantification showing that anterograde SiR-Lys tracks moving through the AIS of DIV7 neurons are significantly fainter compared to those moving anterogradely through the proximal axon of DIV3 neurons. The graph shows individual data points and mean ± SD from 19–25 neurons per condition obtained from three independent cultures. Statistical significance was calculated by two-tailed Welch’s t-test. ***P < 0.001.

**Figure S6.**
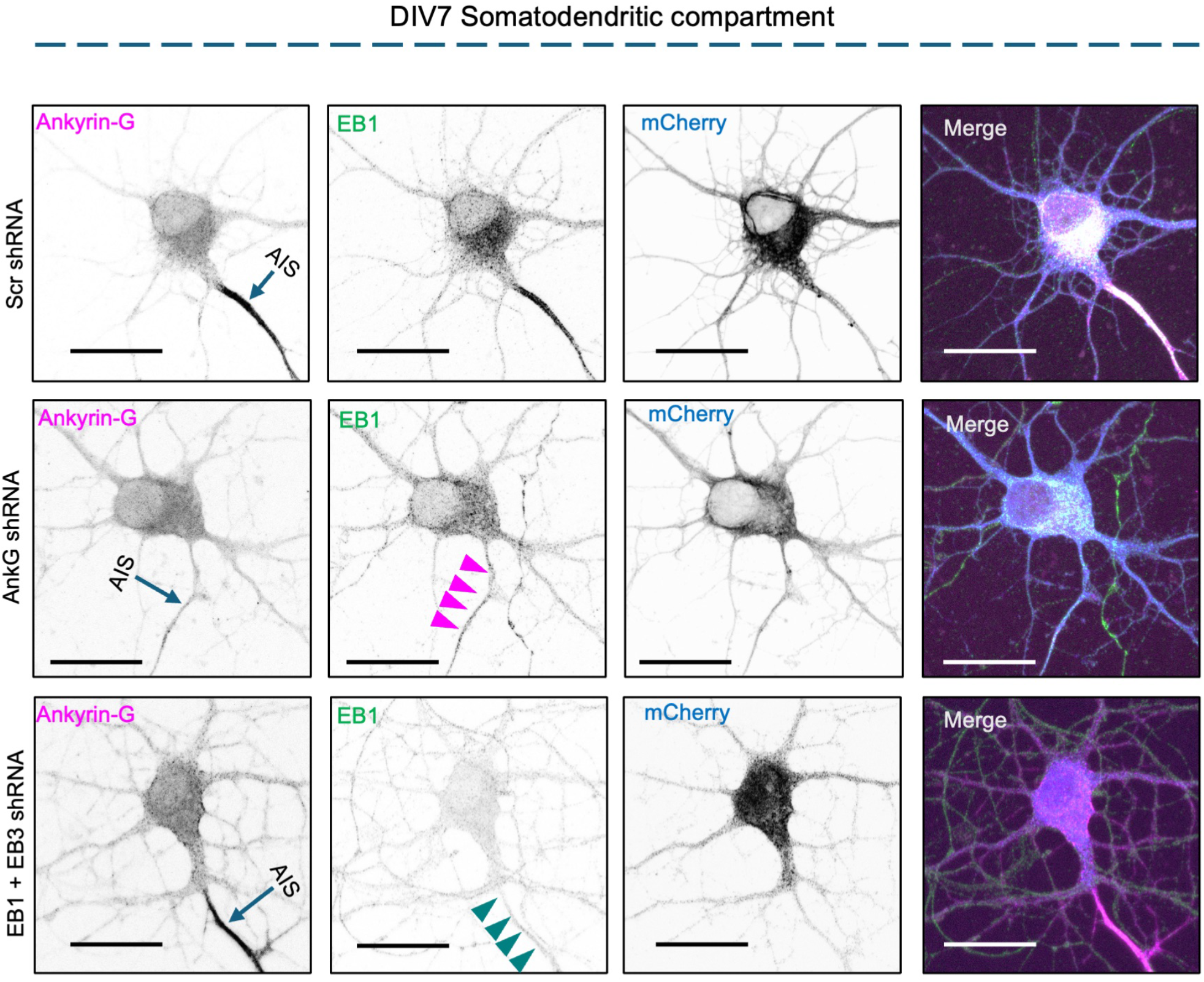
Ankyrin-G and EB1/EB3 depletion differentially alter AIS organization. Confocal fluorescence microscopy of the somatodendritic compartment of DIV7 rat hippocampal neurons expressing Scr shRNA, AnkG shRNA or EB1+EB3 shRNA along with mCherry, fixed, permeabilized, and immunostained for ankyrin-G and EB1. Single-channel images are shown in grayscale, and merged images in color with ankyrin-G in magenta, EB1 in green and mCherry in blue. The AIS is indicated. AnkG depletion reduces AnkG staining and is accompanied by loss of EB1 localization (magenta arrowheads), whereas upon EB1+EB3 depletion discernible ankyrin-G staining remains at the AIS but EB1 staining is markedly reduced (teal arrowheads). Scale bars: 20 μm.

**Figure S7.**
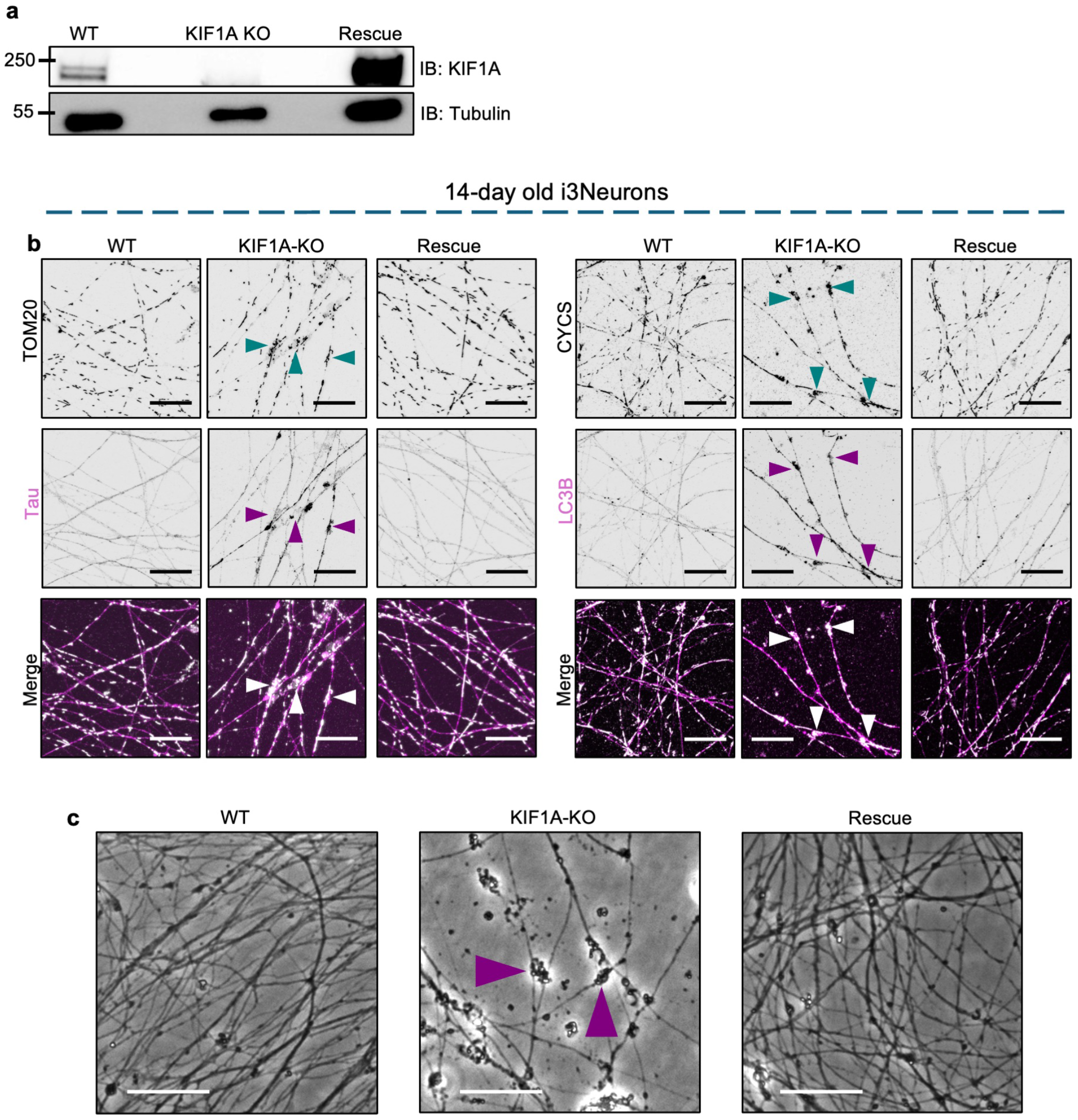
Validation of KIF1A knockout and rescue and associated axonal defects in human i3Neurons. **a,** Immunoblot analysis of lysates from WT, KIF1A-KO, and KIF1A-KO i3Neurons rescued with human KIF1A (Rescue), showing loss of KIF1A expression in KIF1A-KO neurons and restoration following rescue. Tubulin was used as a loading control. Molecular masses are indicated in kilodaltons. **b,** Confocal fluorescence microscopy of primarily axonal fields from 14-day-old WT, KIF1A-KO, and rescued i3Neurons. Left – neurons immunostained for the mitochondrial protein TOM20 and the axonal marker Tau. Single-channel images are shown in grayscale, and merged images in color with TOM20 in white and Tau in magenta. Right – neurons immunostained for cytochrome c (CYCS) and the autophagosome marker LC3B. Single-channel images are shown in grayscale, and merged images in color with CYCS in white and LC3 in magenta. Green arrowheads indicate enlarged axonal regions containing accumulated mitochondria and magenta arrowheads indicate corresponding accumulation of Tau and LC3Β-positive structures in KIF1A-KO neurons. Scale bars: 20 μm. **c,** Representative transmitted-light images of 14-day-old WT, KIF1A-KO, and rescued i3Neurons. Arrowheads indicate prominent neurite swellings in KIF1A-KO neurons that are reduced following KIF1A rescue. Scale bars: 200 μm.

**Supplementary Video 1. mNG-KIF1A and SiR-Lys movement through DIV3 proximal axon** Live-cell spinning-disk fluorescence microscopy of the proximal axon of DIV3 rat hippocampal neurons expressing mNG or mNG-KIF1A and stained with SiR-Lys. mNG or mNG-KIF1A is shown in green and SiR-Lys in magenta. Arrowheads indicate representative anterograde moving vesicles.

**Supplementary Video 2. mNG-KIF1A and SiR-Lys movement through DIV7 AIS** Live-cell spinning-disk fluorescence microscopy of the AIS of DIV7 rat hippocampal neurons expressing mNG or mNG-KIF1A and stained with SiR-Lys. mNG or mNG-KIF1A is shown in green and SiR-Lys in magenta. White arrowheads indicate representative anterograde moving SiR-Lys vesicles and green arrowheads indicate representative anterograde moving mNG-KIF1A puncta not colocalized with SiR-Lys.

**Supplementary Video 3. Comparison of LAMP1-mNG and SiR-Lys co-movement** Live-cell spinning-disk fluorescence microscopy comparing LAMP1-mNG and SiR-Lys movement through the proximal axon of DIV3 neurons and the AIS of DIV7 neurons. LAMP1-mNG is shown in green and SiR-Lys in magenta. White arrowheads indicate representative anterograde moving LAMP1-mNG and SiR-Lys colocalized vesicles and green arrowheads indicate representative anterograde moving LAMP1-mNG puncta not colocalized with SiR-Lys.

**Supplementary Video 4. Effect of AnkG/EB shRNA on mNG-KIF1A and SiR-Lys co-movement** Live-cell spinning-disk fluorescence microscopy of the AIS of DIV7 rat hippocampal neurons expressing Scr shRNA, AnkG shRNA, or EB1+EB3 shRNA together with mNG-KIF1A and stained with SiR-Lys. mNG-KIF1A is shown in green and SiR-Lys in magenta. White arrowheads indicate representative anterograde moving SiR-Lys vesicles and green arrowheads indicate representative anterograde moving mNG-KIF1A puncta not colocalized with SiR-Lys.

**Supplementary Video 5. Effect of T258M mutation on mNG-KIF1A and SiR-Lys co-movement** Live-cell spinning-disk fluorescence microscopy of the AIS of DIV7 rat hippocampal neurons expressing mNG, mNG-KIF1A-WT, or mNG-KIF1A-T258M and stained with SiR-Lys. mNG or mNG- KIF1A is shown in green and SiR-Lys in magenta. Arrowheads indicate representative anterograde moving vesicles.

## Declaration of generative AI and AI-assisted technologies

During the preparation of this work, the authors used ChatGPT (OpenAI) to assist with literature searching, testing Fiji macros, and to check manuscript spelling, grammar and punctuation. The authors reviewed and edited all AI-assisted output and take full responsibility for the content of the article.

